# Microbial Biosynthesis of Lactate Esters

**DOI:** 10.1101/498576

**Authors:** Jong-Won Lee, Cong T. Trinh

## Abstract

**Background:** Green organic solvents such as lactate esters have broad industrial applications and favorable environmental profiles. Thus, manufacturing and use of these biodegradable solvents from renewable feedstocks help benefit the environment. However, to date, the direct microbial biosynthesis of lactate esters from fermentable sugars has not yet been demonstrated.

**Results:** In this study, we present a microbial conversion platform for direct biosynthesis of lactate esters from fermentable sugars. First, we designed a pyruvate-to-lactate ester module, consisting of a lactate dehydrogenase (*ldhA*) to convert pyruvate to lactate, a propionate CoA-transferase (*pct*) to convert lactate to lactyl-CoA, and an alcohol acyltransferase (*AAT*) to condense lactyl-CoA and alcohol(s) to make lactate ester(s). By generating a library of five pyruvate-to-lactate ester modules with divergent AATs, we screened for the best module(s) capable of producing a wide range of linear, branched, and aromatic lactate esters with an external alcohol supply. By co-introducing a pyruvate-to-lactate ester module and an alcohol (i.e., ethanol, isobutanol) module into a modular *Escherichia coli* (chassis) cell, we demonstrated for the first time the microbial biosynthesis of ethyl and isobutyl lactate esters directly from glucose. In an attempt to enhance ethyl lactate production as a proof-of-study, we re-modularized the pathway into 1) the upstream module to generate the ethanol and lactate precursors and 2) the downstream module to generate lactyl-CoA and condense it with ethanol to produce the target ethyl lactate. By manipulating the metabolic fluxes of the upstream and downstream modules through plasmid copy numbers, promoters, ribosome binding sites, and environmental perturbation, we were able to probe and alleviate the metabolic bottlenecks by improving ethyl lactate production by 4.96-fold. We found that AAT is the most rate limiting step in biosynthesis of lactate esters likely due to its low activity and specificity towards the non-natural substrate lactyl-CoA and alcohols.

**Conclusions:** We have successfully established the biosynthesis pathway of lactate esters from fermentable sugars and demonstrated for the first time the direct fermentative production of lactate esters from glucose using an *E. coli* modular cell. This study defines a cornerstone for the microbial production of lactate esters as green solvents from renewable resources with novel industrial applications.

## BACKGROUND

Solvents are widely used as primary components of cleaning agents, adhesives, and coatings and in assisting mass and heat transfer, separation and purification of chemical processes [1]. However, these solvents are volatile organic compounds (VOCs) that contribute to ozone depletion and photochemical smog via free radical air oxidation and hence cause many public health problems such as eye irritation, headache, allergic skin reaction, and cancer [1, 2]. Thus, recent interest in use of alternative green solvents is increasing due to environmental regulation and compelling demand for the eco-friendly solvents derived from renewable sources [3, 4].

Lactate esters are platform chemicals that have a broad range of industrial applications in flavor, fragrance, and pharmaceutical industries [5]. These esters are generally considered as green solvents because of their favorable toxicological and environmental profiles. For instance, ethyl lactate is 100% biodegradable, non-carcinogenic, non-corrosive, low volatile, and unhazardous to human health and the environment [6]. Due to the unique beneficial properties of ethyl lactate, it has been approved as a Significant New Alternatives Policy (SNAP) solvent by the U.S. Environmental Protection Agency (EPA) and as food additives by the U.S. Food and Drug Administration (FDA) [6]. Recent technical and economic analysis conducted by the National Renewable Energy Laboratory (NREL) considers ethyl lactate to be one of the top twelve bioproducts [7].

In industrial chemical processes, lactate esters are currently produced by esterification of lactic acid with alcohols using homogenous catalysts (e.g., sulfuric acid, hydrogen chloride, and/or phosphoric acid) under high temperature reaction conditions [8]. However, use of strong acids as catalysts cause corrosive problems and often require more costly equipment for process operation and safety. Furthermore, the esterification reactions are thermodynamically unfavorable (Δ*G* = +5 kcal/mol) in aqueous solutions and often encounter significant challenge due to self-polymerization of lactate [9]. Alternatively, microbial catalysts can be harnessed to produce these esters from renewable and sustainable feedstocks in a thermodynamically favorable reaction (Δ*G* = −7.5 kcal/mol) in an aqueous phase environment at room temperature and atmospheric pressure [10–16]. This reaction uses an alcohol acyltransferase (AAT) to generate an ester by condensing an alcohol and an acyl-CoA. AAT can catalyze a broad substrate range including i) linear or branched short-to-long chain fatty alcohols [10, 11, 17], ii) aromatic alcohols [18], and iii) terpenols [19–22] as well as various fatty acyl-CoAs [11, 13]. To date, while microbial biosynthesis of the precursor metabolites for lactate esters have been well established such as lactate [13, 16, 23–27], lactyl-CoA [28–30], ethanol [31, 32], propanol [33], isopropanol [34], butanol [35], isobutanol [36], amyl alcohol [37], isoamyl alcohol [38], benzyl alcohol [39], 2-phenylethanol [40, 41], and terpenols [19–22], the direct microbial biosynthesis of lactate esters from fermentable sugars has not yet been demonstrated.

In this work, we aimed to demonstrate the feasibility of microbial production of lactate esters as green organic solvents from renewable resources. To enable the direct microbial biosynthesis of lactate esters from fermentable sugars, we first screened for an efficient AAT suitable for lactate ester production using a library of five pyruvate-to-lactate ester modules with divergent AATs. We next demonstrated direct fermentative biosynthesis of ethyl and isobutyl lactate esters from glucose by co-introducing a pyruvate-to-lactate ester module and an alcohol module (i.e., ethanol and isobutanol) into an engineered *Escherichia coli* modular cell. As a proof-of-study to improve ethyl lactate production, we employed a combination of metabolic engineering and synthetic biology approaches to dissect the pathway to probe and alleviate the potential metabolic bottlenecks.

## RESULTS AND DISCUSSION

### *In vivo* screening of efficient AATs critical for lactate ester biosynthesis

The substrate specificity of AATs is critical to produce target esters [13]. For example, ATF1 exhibits substrate preference for biosynthesis of acyl (C4-C6) acetates while SAAT and VAAT prefer biosynthesis of ethyl (C2-C6) acylates. Even though both SAAT and VAAT are derived from the same strawberry genus, they also show very distinct substrate preferences; specifically, SAAT prefers longer (C4-C6) acyl-CoAs whereas VAAT prefers shorter (C2-C4) acyl-CoAs. To date, none of AATs have been tested for lactate ester biosynthesis. Thus, to enable lactate ester biosynthesis, we began with identification of the best AAT candidate. We designed, constructed, and characterized a library of five pyruvate-to-lactate ester modules (pJW002-006) carrying five divergent AATs including ATF1, ATF2, SAAT, VAAT, and AtfA, respectively. AtfA was used as a negative control because it prefers long-chain acyl-CoAs (C14-C18) and alcohols (C14-C18) [42]. For characterization, 2 g/L of ethanol, propanol, butanol, isobutanol, isoamyl alcohol, and benzyl alcohol were added to culture media with 0.5 mM of IPTG for pathway induction to evaluate biosynthesis of six different lactate esters including ethyl lactate, propyl lactate, butyl lactate, isobutyl lactate, isoamyl lactate, and benzyl lactate, respectively, in high-cell density cultures (Fig. 1A).

**Figure 1.**
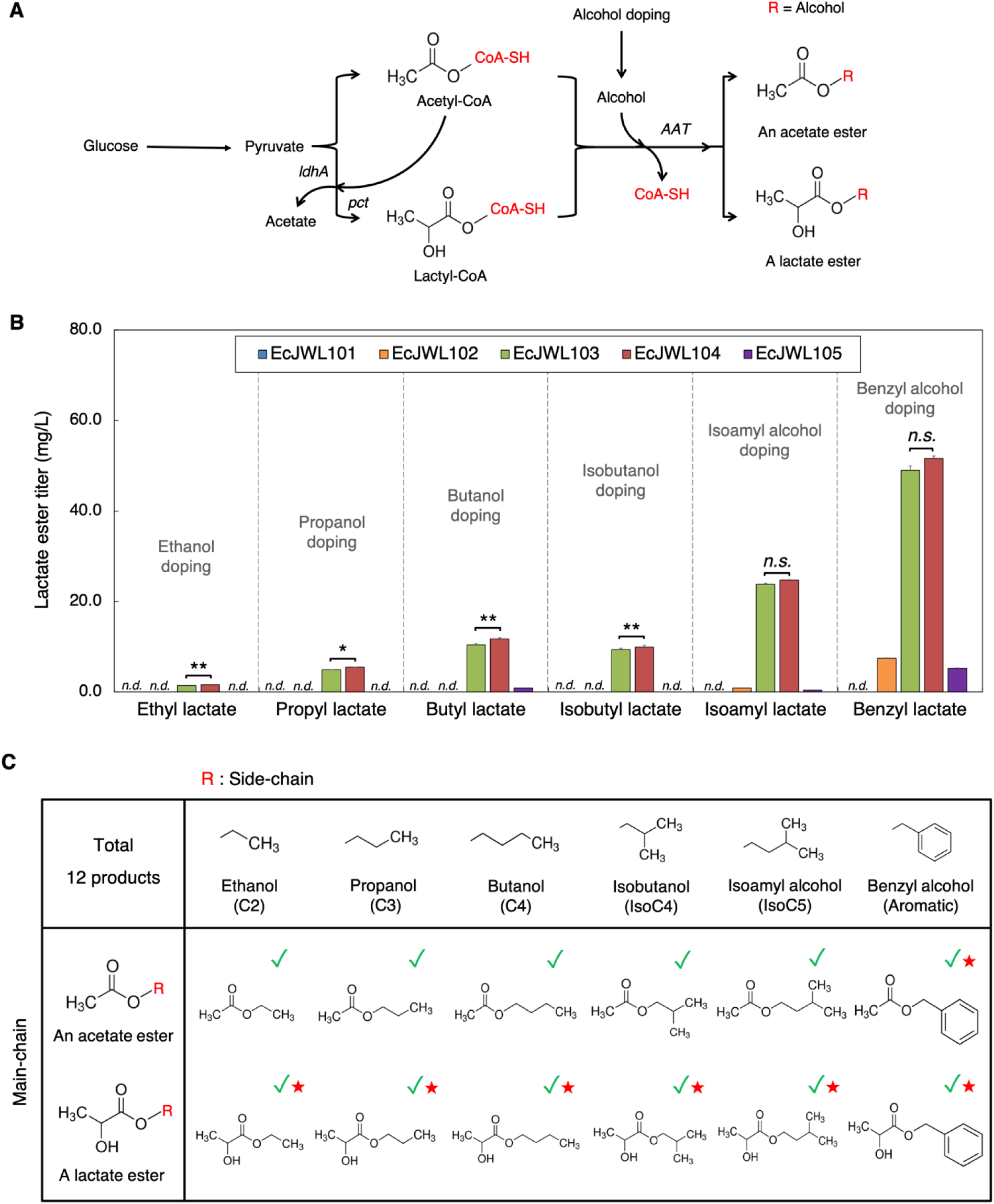
*In vivo* characterization of various alcohol acyltransferases for biosynthesis of lactate esters. **(A)** Biosynthesis pathways of lactate and acetate esters with external supply of alcohols. **(B)** Ester production of EcJW101, EcJW102, EcJW103, EcJW104, and EcJW105 harboring *ATF1*, *ATF2*, *SAAT*, *VAAT*, and *atfA*, respectively in high cell density cultures with various alcohol doping. Each error bar represents 1 standard deviation (s.d., *n=3*). Symbols: *n.s.*, not significant, \**p*-value < 0.073, and \*\**p*-value < 0.013 (Student’s test). **(C)** The library of esters produced. Green check marks indicate the esters produced in this study while red star marks indicate the esters produced for first time in engineered strains.

The results show that most of the strains could produce different types of lactate esters with external supply of alcohols (Fig. 1B, 1C). EcJW104 achieved the highest titer of lactate esters in all cases, producing 1.59 ± 0.04 mg/L of ethyl lactate, 5.46 ± 0.25 mg/L of propyl lactate, 11.75 ± 0.43 mg/L of butyl lactate, 9.92 ± 0.08 mg/L of isobutyl lactate, 24.73 ± 0.58 mg/L of isoamyl lactate, and 51.59 ± 2.09 mg/L of benzyl lactate in ethanol, propanol, butanol, isobutanol, isoamyl alcohol, and benzyl alcohol doping, respectively. The lactate ester biosynthesis of EcJW104 exhibited different alcohol substrate preference in the following order: benzyl alcohol > isoamyl alcohol > butanol > isobutanol > propanol > ethanol (Fig. 1B, Supplementary Table S2).

Due to the presence of endogenous acetyl-CoA, we also produced acetate esters in addition to lactate esters (Fig. 1). Among the strains, EcJW101 achieved the highest titers of acetate esters in all cases, producing 115.52 ± 4.83 mg/L of ethyl acetate, 801.62 ± 33.51 mg/L of propyl acetate, 1,017.90 ± 20.21 mg/L of butyl acetate, 1,210.40 ± 24.83 mg/L of isobutyl acetate, 692.73 ± 7.65 mg/L of isoamyl acetate, and 1,177.98 ± 45.72 mg/L of benzyl acetate in ethanol, propanol, butanol, isobutanol, isoamyl alcohol, and benzyl alcohol doping, respectively. EcJW101 showed different alcohol substrate preference for the acetate ester biosynthesis in the following order: isobutanol > benzyl alcohol > butanol > propanol > isoamyl alcohol > ethanol (Supplementary Table S2).

Taken altogether, VAAT and ATF1 are the most suitable AATs for biosynthesis of lactate esters and acetate esters, respectively. Among the library of 12 esters (Fig. 1C), seven of these esters, including ethyl lactate, propyl lactate, butyl lactate, isobutyl lactate, isoamyl lactate, benzyl lactate, and benzyl acetate, were demonstrated for *in vivo* production in microbes for the first time. EcJW104 that harbors the pyruvate-to-lactate module with *VAAT* could produce 6 out of 6 target lactate esters including ethyl, propyl, butyl, isobutyl, isoamyl, and benzyl lactate. Since EcJW104 achieved the highest titer of lactate esters in all cases, it was selected for establishing the biosynthesis pathway of lactate esters from glucose.

### Establishing the lactate ester biosynthesis pathways

We next demonstrated direct fermentative production of lactate esters from glucose using the best VAAT candidate. We focused on the biosynthesis of ethyl and isobutyl lactate esters. We designed the biosynthesis pathways for ethyl and isobutyl lactate by combining the pyruvate-to-lactate ester module (pJW005) with the ethanol (pCT24) and isobutanol (pCT13) modules, respectively. By co-transforming pJW005/pCT24 and pJW005/pCT13 into the modular cell EcDL002, we generated the production strains, EcJW201 and EcJW202, for evaluating direct conversion of glucose to ethyl and isobutyl lactate esters.

We characterized EcJW201 and EcJW202 together with the parent strain, EcDL002, as a negative control in high-cell density cultures. The results show EcJW201 and EcJW202 produced ethyl (Fig. 2A) and isobutyl (Fig. 2B) lactate from glucose, respectively, while the negative control strain EcDL002 could not. Consistently, the expressions of metabolic enzymes of the ethyl and isobutyl lactate pathways were confirmed in EcJW201 and EcJW202, respectively, by SDS-PAGE analysis (Supplementary Figure S1). During 24 h fermentation, EcJW201 produced 2.24 ± 0.28 mg/L of ethyl lactate with a specific productivity of 0.04 ± 0.00 mg/gDCW/h while EcJW202 produced 0.26 ± 0.01 mg/L of isobutyl lactate with a specific productivity of 0.01 ± 0.00 mg/gDCW/h. In addition to ethyl or isobutyl lactate biosynthesis, EcJW201 also produced 92.25 ± 9.20 mg/L of ethyl acetate while EcJW202 generated 1.36 ± 0.74 mg/L of ethyl acetate and 0.34 ± 0.07 mg/L of isobutyl acetate (Supplementary Table S3A). Taken altogether, the direct microbial synthesis of lactate esters from fermentable sugar was successfully demonstrated. Since the lactate ester production was low, the next logical step was to identify and alleviate the key pathway bottlenecks for enhanced lactate ester biosynthesis. As proof-of-principle, we focused on optimization of the ethyl lactate production as presented in the subsequent sections.

**Figure 2.**
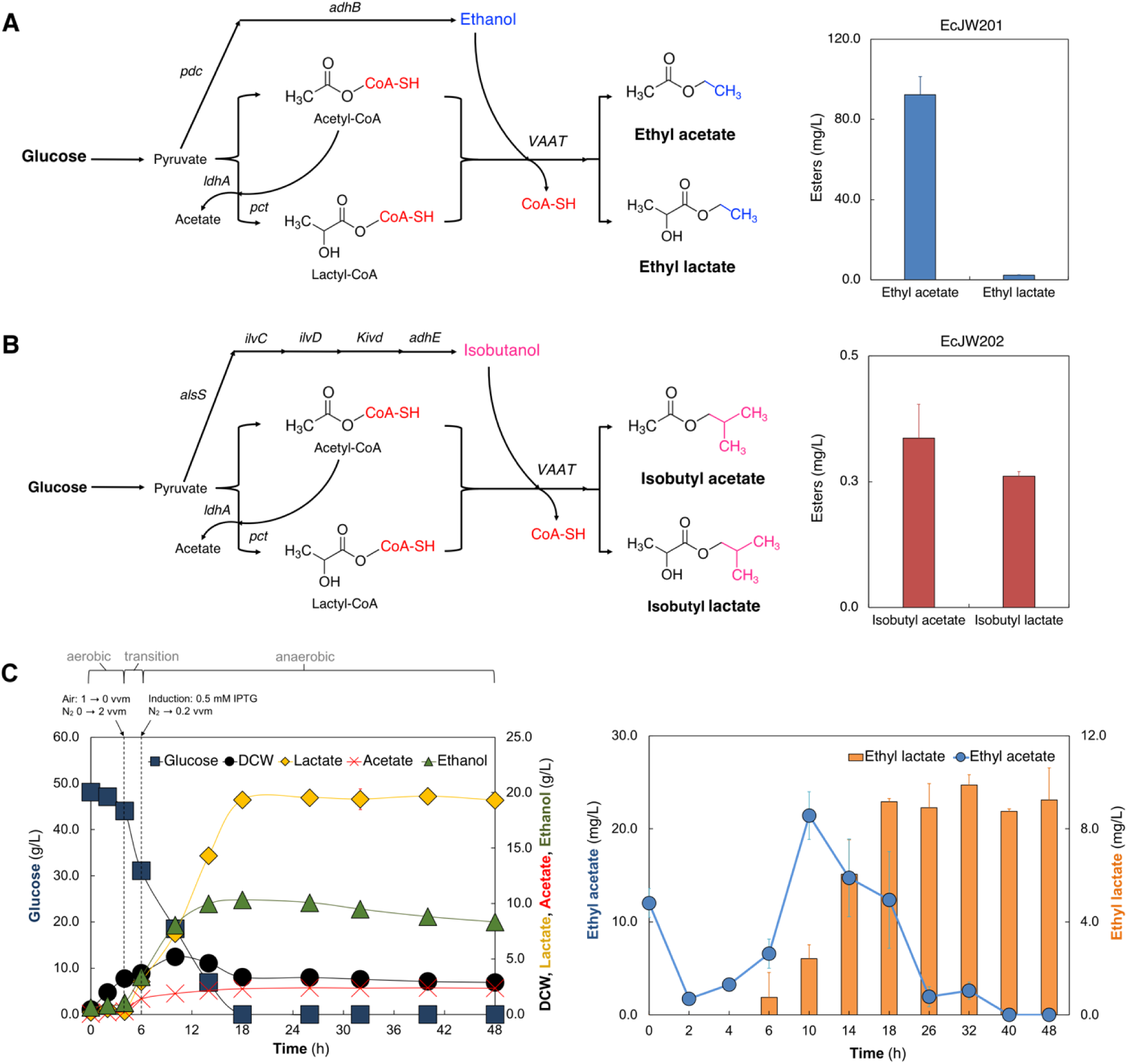
Design, construction, and validation of the lactate ester biosynthesis pathways in *E. coli*. **(A)** Engineered biosynthesis pathway of ethyl lactate from glucose and its production in high cell density culture of EcJW201. **(B)** Engineered biosynthesis pathway of isobutyl lactate from glucose and its production in high cell density culture of EcJW202. In Fig. 2A and 2B, all of the strains were induced at 0 h with 0.5 mM IPTG. Each error bar represents 1 s.d. (*n=3*). **(C)** Production of ethyl lactate from glucose in pH-controlled batch fermentation of EcJW201. The strain was induced at 6 h with 0.5 mM IPTG. Each error bar represents 1 s.d. (*n=2*).

### Identifying and alleviating key bottlenecks of the ethyl lactate biosynthesis pathway

#### Evaluating the biosynthesis of ethyl lactate in pH-controlled fermentation as a basis to identify potential pathway bottlenecks

In an attempt to identify the key bottlenecks of the ethyl lactate biosynthesis pathway, we characterized EcJW201 in pH-controlled bioreactors. The results show that EcJW201 produced 9.17 ± 0.12 mg/L of ethyl lactate with a specific productivity of 0.15 ± 0.02 mg/gDCW/h and a yield of 0.19 ± 0.00 mg/g glucose (Fig. 2C, Supplementary Table S3B) in 18 h. Under pH-controlled fermentation, EcJW201 achieved 4.09-fold (from 2.24 ± 0.28 to 9.17 ± 0.12 mg/L), 3.75-fold (from 0.04 ± 0.00 to 0.15 ± 0.02 mg/gDCW/h), and 19-fold (from 0.01 ± 0.00 to 0.19 ± 0.00 mg/g glucose) improvement in titer, specific productivity, and yield, respectively, as compared to the strain performance in the high cell density culture. It is interesting to observe that ethyl acetate was first produced then consumed after 10 h, which is likely due to the endogenous esterase of *E. coli* as observed in a recent study [43]. Different from ethyl acetate, we did not observe ethyl lactate degradation during fermentation, especially after glucose was depleted. Even though the strain performance in pH-controlled bioreactors was enhanced by increased supply of precursor metabolites (19.35±0.29 g/L of lactate and 10.31± 0.41 g/L of ethanol, Supplementary Table S3B) from higher concentration of carbon source, the titer of ethyl lactate did not increase during the fermentation. This result suggests that (i) rate-limiting conversion of lactate into lactyl-CoA by Pct and/or condensation of lactyl-CoA with an ethanol by VAAT and/or (ii) toxicity of ethyl lactate on *E. coli* health might have limited lactate ester biosynthesis. Therefore, to enhance ethyl lactate production, it is important to elucidate and alleviate these identified potential bottlenecks.

#### Ethyl lactate exhibited minimal toxicity on cell growth among lactate esters

To determine whether lactate esters inhibited cell growth and hence contributed to low lactate ester production, we cultured the parent strain, EcDL002, in a microplate reader with or without supply of various concentrations of lactate esters including ethyl, propyl, butyl, isobutyl, isoamyl, or benzyl lactate. The results show that ethyl lactate was the least toxic among the six lactate esters characterized where the growth rate (0.47 ± 0.04 1/h) and cell titer (OD = 0.42 ± 0.03) decreased by 6% and 10%, respectively, upon cell exposure to 5 g/L ethyl lactate. On the other hand, isoamyl lactate was the most toxic among the lactate esters, where cell exposure to only 0.5 g/L ester resulted in 18% and 15% reduction in the growth rate (0.41 ± 0.02 1/h) and OD (0.40 ± 0.03), respectively (Supplementary Figure S2A). The toxicity of lactate esters can be ranked in the following order: isoamyl lactate > benzyl lactate > butyl lactate > isobutyl lactate > propyl lactate > ethyl lactate. There existed a positive correlation between the logP values of lactate esters and their degrees of toxicity (Supplementary Figure S2B). This result was consistent with literature, illustrating that increasing toxicity of esters is highly correlated with increasing chain length of alcohol moieties that can severely disrupt cell membrane [44]. It should be note that since *E. coli* can effectively secrete short-chain esters [10], external exposure of cells to lactate esters in our experiment design is sufficient to probe the potential toxicity caused by endogenous production of these esters. Taken altogether, ethyl lactate is the least toxic and was not likely the main reason for the low production of ethyl lactate observed. It was likely the downstream pathway, responsible for conversion of lactate into lactyl-CoA by Pct and/or condensation of lactyl-CoA with ethanol by VAAT, might have been contributed to the inefficient ethyl lactate biosynthesis.

#### Downstream pathway of the lactate ester biosynthesis is the key bottleneck

To identify and alleviate the ethyl lactate biosynthesis pathway, we re-modularized it with two new parts: i) the upstream module carrying *ldhA*, *pdc*, and *adhB* for production of lactate and ethanol from sugar and ii) the downstream module carrying *pct* and *VAAT* for converting lactate into lactyl-CoA and condensing lactyl-CoA and ethanol (Fig. 3A). We controlled metabolic fluxes of these modules by manipulating their plasmid copy numbers and levels of promoter induction with IPTG. By introducing the plasmids pJW007-015 into EcDL002, we generated the strains EcJW106-108 and EcJW203-208, respectively (Fig. 3B). To evaluate the performance of these constructed strains for ethyl lactate production, we characterized them in high cell density cultures induced with various concentrations of IPTG (0.01, 0.1, and 1.0 mM).

**Figure 3.**
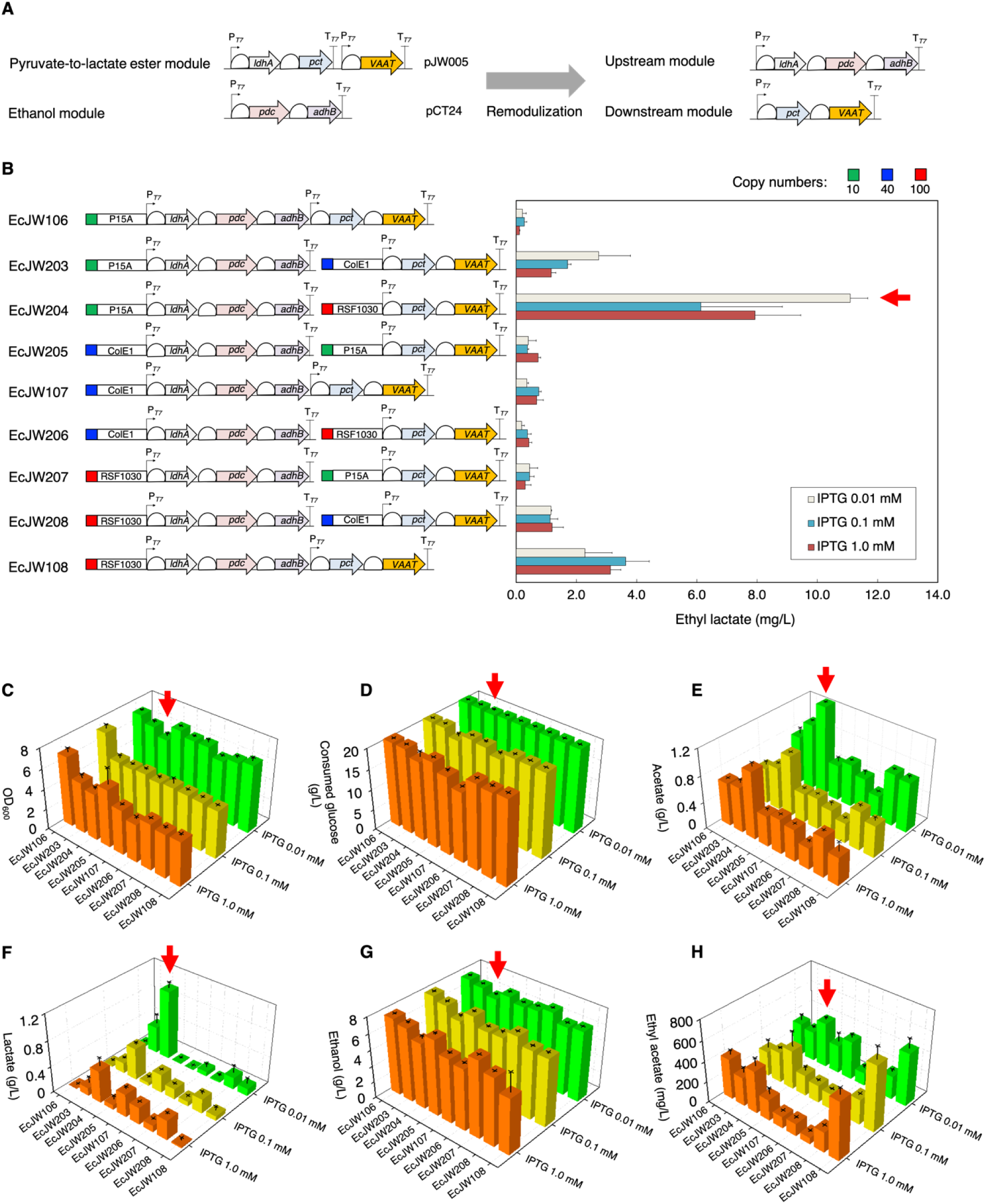
Combinatorial modular pathway optimization of enhanced ethyl lactate biosynthesis by varying plasmid copy number. **(A)** Re-modularization of the ethyl lactate biosynthesis pathway. Pyruvate-to-lactate ester and ethanol modules were re-modulated into upstream and downstream modules using plasmids with different copy numbers. **(B)** Ethyl lactate production, **(C)** OD_600_, **(D)** consumed glucose, **(E)** acetate, **(F)** lactate, **(G)** ethanol, and **(H)** ethyl acetate of EcJW106-108 and EcJW203-208 in high cell density cultures induced with various concentrations of IPTG. Green rectangle: low copy number plasmid (10); P15A: origin of pACYCDuet-1; Blue rectangle: medium copy number plasmid (40); ColE1: origin of pETDuet-1; Red rectangle: high copy number plasmid (100); RSF1030: origin of pRSFDuet-1; P_T7_: T7 promoter; T_T7_: T7 terminator. All of the strains were induced at 0 h with 0.01, 0.1, or 1.0 mM IPTG, respectively. Each error bar represents 1 s.d. (*n=3*). Red arrows indicate the selected strain with an optimum concentration of IPTG for the further studies.

The results show that EcJW204, carrying the upstream module with a low copy number plasmid (P15A origin) and the downstream module with a high copy number plasmid (RSF1030 origin) induced by 0.01 mM of IPTG, achieved the highest titer of ethyl lactate. As compared to EcJW201, EcJW204 achieved 4.96-fold (an increase from 2.24 to 11.10 ± 0.58 mg/L), 5.50-fold (from 0.04 ± 0.00 to 0.22 ± 0.02 mg/gDCW/h), and 54.0-fold (from 0.01 ± 0.00 to 0.54 ± 0.04 mg/g glucose) improvement in titer, specific productivity, and yield of ethyl lactate, respectively (Fig. 3B, Supplementary Table S5). Upon IPTG induction at 24 h, we observed the reduced cell growth of the host strains with use of high concentration of IPTG (Fig. 3C, Supplementary Table S4), suggesting that they suffered from metabolic burden due to overexpression of multiple enzymes [45] and also explaining why use of low concentration of IPTG can help yield better production of ethyl lactate.

Although EcJW204 showed better performance in ethyl lactate production than EcJW201, the accumulation of lactate and ethanol was still observed (Fig. 3F and 3G, Supplementary Table S4), indicating the pathway bottleneck remained. In particular, the downstream module flux was outcompeted by the upstream module flux and hence failed to turn over the precursor metabolites quickly enough. This result helps explain why a combination of the upstream module (for producing lactate and ethanol from sugar) with a low copy number plasmid and the downstream module (for converting lactate into lactyl-CoA and condensing lactyl-CoA and ethanol) with a high copy number plasmid outperformed eight other combinations. Notably, the best ethyl lactate producer EcJW204 achieved the highest lactate and lowest ethanol production among the nine characterized strains (Fig. 3F and 3G, Supplementary Table S4), suggesting redistribution of the carbon flux from ethanol to lactate likely helped improve ethyl lactate production. Thus, we hypothesized that redistribution of the carbon source from ethanol to lactate would help to improve ethyl lactate production. To test this hypothesis, we first examined whether i) downregulation of the ethanol flux of the upstream module enabled redistribution of the carbon flow from ethanol to lactate and ii) this redistribution could improve ethyl lactate production before proceeding to investigate the potential bottleneck of downstream module.

#### High ethanol synthesis of the upstream module was critical for ethyl lactate biosynthesis due to low specificity and activity of AAT

To downregulate the ethanol flux of the upstream module, we first reconfigured pJW007, the upstream module of the best performer EcJW204, with a library of two weaker promoters and four weaker synthetic RBSs (Fig. 4A, Supplementary Figure S3A), resulting in four new upstream modules (pJW019-022). By introducing each newly constructed upstream module into EcDL002 together with the downstream module pJW012 used in EcJW204, we next generated the strains EcJW209-212 and characterized them in high cell density cultures induced with 0.01 mM IPTG.

**Figure 4.**
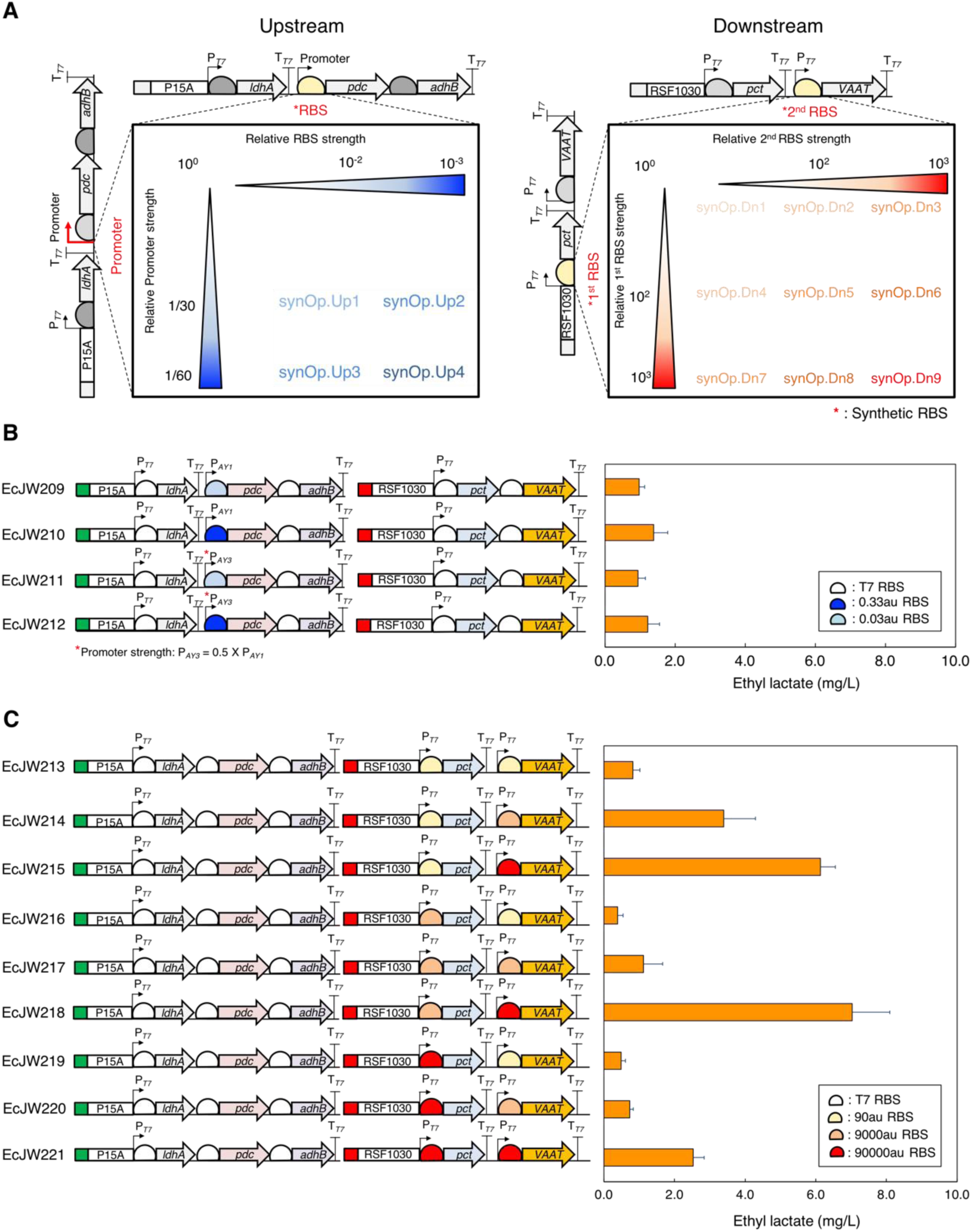
Probing and alleviating the potential metabolic bottlenecks of the upstream or downstream modules of EcJW204 by varying the strength of promoters and/or ribosome binding sites. **(A)** Design of synthetic operons for the upstream and downstream modules. For the upstream module, the T7 promoter in MCS2 and the RBS between T7 promoter in MCS2 and the start codon of *pdc* were replaced with the combination of P_AY1_ or P_AY3_ promoter and 0.3 or 0.03au RBS. For the downstream module, the RBS between T7 promoter in MCS1 and the start codon of *pct* gene and the RBS between T7 promoter in MCS2 and the start codon of *VAAT* gene were replaced with the combination of 90, 9000, or 90000au RBS and 90, 9000, or 90000au RBS, respectively. Production of ethyl lactate in high cell density cultures of **(B)** EcJW209-212 and **(C)** EcJW213-221. Green rectangle: low copy number plasmid (10); P15A: origin of pACYCDuet-1; Red rectangle: high copy number plasmid (100); RSF1030: origin of pRSFDuet-1; P_T7_: T7 promoter; T_T7_: T7 terminator. All of the strains were induced at 0 h with 0.01 mM IPTG. Each error bar represents 1 s.d. (*n=3*).

The results show that while the carbon flux was successfully redistributed from ethanol to lactate, resulting in 5.97∼6.92-fold decrease in ethanol production (from 8.30 ± 0.17 to 1.39 ± 0.10 ∼ 1.20 ± 0.01 g/L) and 1.67∼2.59-fold increase in lactate production (from 1.06 ± 0.09 to 1.77 ± 0.37 g/L∼2.75 ± 0.09 g/L) (Supplementary Table S6A), the ethyl lactate production was reduced by 7.99∼11.81-fold in ethyl lactate production (from 11.10 ± 0.58 to 1.39 ± 0.40 ∼ 0.94 ± 0.22 mg/L) in all four characterized strains as compared to that of EcJW204 (Fig. 4B, Supplementary Table S6B). This result suggests that a high level of ethanol is critical for VAAT to produce ethyl lactate.

To support this conclusion, we evaluated the effect of external ethanol supply on production of ethyl esters in high cell density cultures of EcJW209-212 induced with 0.01 mM IPTG. Indeed, with external ethanol supply, we observed enhanced production of both ethyl lactate and ethyl acetate in EcJW209-212. Specifically, with addition of 2 g/L of ethanol, the ethyl lactate and ethyl acetate production increased by 2.27 ∼ 3.33-fold (from 1.39 ± 0.40 to 3.15 ± 0.15 mg/L ∼ from 0.98 ± 0.15 to 3.26 ± 0.26 mg/L) and 1.27∼2.07-fold (from 36.46 ± 3.86 to 46.22 ± 1.33 mg/L ∼ from 21.96 ± 0.84 to 45.40 ± 1.20 mg/L), respectively (Supplementary Table S6). Further addition of ethanol up to 10 g/L improved the ethyl lactate and ethyl acetate production by 3.78∼5.26-fold (from 1.39 ± 0.40 to 5.26 ± 0.27 mg/L ∼ from 0.94 ± 0.15 mg/L to 4.49 ± 0.41 mg/L) and 4.09∼6.92-fold (from 36.46 ± 3.86 to 148.97 ± 3.80 mg/L ∼ from 21.96 ± 0.84 mg/L to 151.87 ± 2.34 mg/L), respectively (Supplementary Table S6). Interestingly, while the total titer of ethyl esters increased with the increasing addition of ethanol (Fig. 5A), the proportion of ethyl lactate in the total ester slightly increased in the range of 3.2∼7.0% (Fig. 5B), suggesting that VAAT prefers acetyl-CoA over lactyl-CoA with ethanol as a co-substrate. Notably, we observed a strong linear correlation between ethyl esters production and the amount of added ethanol (i.e., for ethyl lactate, R^2^ = 0.85∼0.94; for ethyl acetate, R^2^ = 0.99∼1.00) (Supplementary Figure S4A). The results revealed that abundant availability of ethanol is essential to achieve high production of ethyl esters, indicating the main reason for the improved ethyl lactate production in EcJW204 was most likely due to the upregulation of downstream module with a high copy number plasmid.

**Figure 5.**
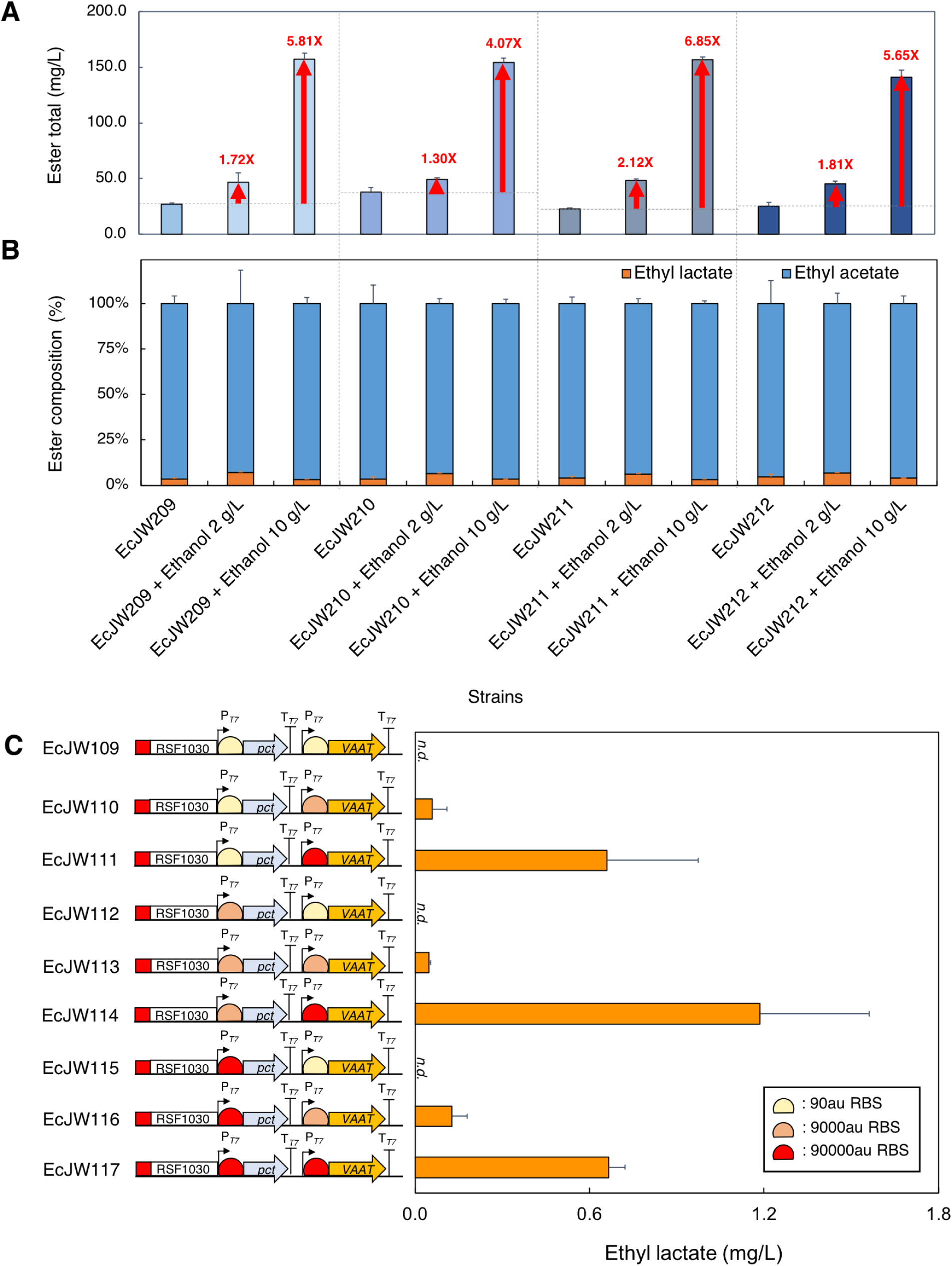
**(A)** Total esters and **(B)** composition of total esters produced in high cell density cultures of EcJW209-212 with or without addition of ethanol. **(C)** Ethyl lactate production of EcJW109-117 with addition of 2 g/L of lactate and ethanol. Red rectangle: high copy number plasmid (100); RSF1030: origin of pRSFDuet-1; P_T7_: T7 promoter; T_T7_: T7 terminator. All of the strains were induced at 0 h with 0.01 mM IPTG. Each error bar represents 1 s.d. (*n=3*).

#### AAT was the most rate limiting step of the downstream module

To determine whether Pct for conversion of lactate to lactyl-CoA or VAAT for condensation of lactyl-CoA and an alcohol was the most rate limiting step of the downstream module, we redesigned and constructed nine downstream modules (pJW027-035) derived from pJW012 of the best performer EcJW204 using a combination of three synthetic RBSs for Pct expression (synRBS_pct#1-3_) and three synthetic RBSs for VAAT expression (synRBS_VAAT#1-3_) (Fig. 4A, Supplementary Figure S3B). We introduced each newly constructed downstream module into EcDL002 together with the original upstream module (pJW007) used in EcJW204 to generate EcJW213-221. Then, we characterized the constructed strains in high cell density cultures induced with 0.01 mM IPTG.

The results show that the strains harboring the stronger RBSs for VAAT expression achieved the higher titers of ethyl lactate and ethyl acetate regardless of the RBS strengths for Pct expression (Fig. 4C, Supplementary Table S7). There is a strong linear correlation between ethyl ester production and the strength of RBS for VAAT expression (Supplementary Figure S4B). To further validate these results without the influence of the upstream module, we additionally constructed the strains EcJW109-117 by introducing nine individual downstream modules (pJW027-035) into EcDL002 and then characterized these strains in high cell density cultures with addition of 2 g/L of lactate, 2 g/L of ethanol, and 0.01 mM of IPTG. We could observe the same strong linear correlation between ethyl ester production and high VAAT expression without the upstream module (Fig. 5C).

Taken altogether, these results suggest that VAAT not Pct was the most rate limiting step of the downstream module of the ethyl lactate biosynthesis pathway. Specifically, a combination of low affinity towards lactyl-CoA and ethanol of VAAT contributed to low ethyl lactate biosynthesis. Further studies on discovery of novel AATs, exhibiting high activity towards lactyl-CoA and alcohols but not acetyl-CoA, together with rational protein engineering of these enzymes would be warranted for improving lactate ester production.

In principle, the lactate ester platform can be controlled to produce enantiomers with broad industrial applications. Since the endogenous *E. coli* D-lactate dehydrogenase (LdhA) was overexpressed in the *ldhA*-deficient modular cell of our study, it is anticipated that D-(-)-lactate and the associated D-(-)-lactate esters were mainly produced. To date, production of optically pure D-(-)-[23] and L-(+)-form [26] of lactate from glucose in *E. coli* [25] has been well established. In addition, *pct* from *C. propionicum* [28] and *Megasphaera elsdenii* [29, 30] have been used for converting D-(-)-lactate into D-(-)-lactyl-CoA in polylactic acid (PLA) production in *E. coli* and their catalytic activity towards L-(+)-lactate has also been demonstrated [46, 47]. Thus, by combining stereospecific Ldh and Pct enzymes together with AATs, it is highly feasible to extend our lactate ester platform for microbial production of stereospecific lactate esters from renewable resources.

## CONCLUSIONS

In this study, we have successfully developed a microbial lactate ester production platform and demonstrated for the first time the microbial biosynthesis of lactate esters directly from fermentable sugars in an *E. coli* modular cell. This study defines a cornerstone for the microbial production of lactate esters as green solvents from renewable resources with novel industrial applications.

## METHODS

### Strain construction

The list of strains used in this study are presented in Table 1. For molecular cloning, *E. coli* TOP10 strain was used. To generate the lactate ester production strains, the modules, including i) the pyruvate-to-lactate ester modules (pJW002-006), ii) the upstream and/or downstream modules (pJW007-pJW028), and iii) the alcohol modules (pCT24 or pCT13), were transformed into the engineered modular *E. coli* chassis cell, EcDL002 [10] via electroporation [48].

**Table 1.** A list of strains used in this study

| Strains | Genotypes | Sources |
| --- | --- | --- |
| *E. coli* TOP10 | F- *mcrA* $\Delta$(*mrr-hsdRMS-mcrBC*) Φ80*lacZ*ΔM15 Δ*lacX74* *recA1* *araD139* $\Delta$(*ara leu*) 7697 *galU galK rpsL* (StrR) *endA1 nupG* | Invitrogen |
| *E. coli* MG1655 | F<sup>-</sup> λ<sup>-</sup> | ATCC 47076 |
| *Clostridium propionicum* | Wildtype | ATCC 25522 |
| EcDL002 | TCS083 (λDE3) Δ*fadE* | [10] |
| EcJW101 | EcDL002/pJW002; amp<sup>R</sup> | This study |
| EcJW102 | EcDL002/pJW003; amp<sup>R</sup> | This study |
| EcJW103 | EcDL002/pJW004; amp<sup>R</sup> | This study |
| EcJW104 | EcDL002/pJW005; amp<sup>R</sup> | This study |
| EcJW105 | EcDL002/pJW006; amp<sup>R</sup> | This study |
| EcJW201 | EcDL002/pJW005 pCT24; amp<sup>R</sup> kan<sup>R</sup> | This study |
| EcJW202 | EcDL002/pJW005 pCT13; amp<sup>R</sup> kan<sup>R</sup> | This study |
| EcJW106 | EcDL002/pJW013; cm<sup>R</sup> | This study |
| EcJW203 | EcDL002/pJW007 pJW011; cm<sup>R</sup> amp<sup>R</sup> | This study |
| EcJW204 | EcDL002/pJW007 pJW012; cm<sup>R</sup> kan<sup>R</sup> | This study |
| EcJW205 | EcDL002/pJW008 pJW010; cm<sup>R</sup> amp<sup>R</sup> | This study |
| EcJW107 | EcDL002/pJW014; amp<sup>R</sup> | This study |
| EcJW206 | EcDL002/pJW008 pJW012; amp<sup>R</sup> kan<sup>R</sup> | This study |
| EcJW207 | EcDL002/pJW009 pJW010; cm<sup>R</sup> kan<sup>R</sup> | This study |
| EcJW208 | EcDL002/pJW009 pJW011; amp<sup>R</sup> kan<sup>R</sup> | This study |
| EcJW108 | EcDL002/pJW015; kan<sup>R</sup> | This study |
| EcJW209 | EcDL002/pJW019 pJW012; cm<sup>R</sup> kan<sup>R</sup> | This study |
| EcJW210 | EcDL002/pJW020 pJW012; cm<sup>R</sup> kan<sup>R</sup> | This study |
| EcJW211 | EcDL002/pJW021 pJW012; cm<sup>R</sup> kan<sup>R</sup> | This study |
| EcJW212 | EcDL002/pJW022 pJW012; cm<sup>R</sup> kan<sup>R</sup> | This study |
| EcJW213 | EcDL002/pJW007 pJW027; cm<sup>R</sup> kan<sup>R</sup> | This study |
| EcJW214 | EcDL002/pJW007 pJW028; cm<sup>R</sup> kan<sup>R</sup> | This study |
| EcJW215 | EcDL002/pJW007 pJW029; cm<sup>R</sup> kan<sup>R</sup> | This study |
| EcJW216 | EcDL002/pJW007 pJW030; cm<sup>R</sup> kan<sup>R</sup> | This study |
| EcJW217 | EcDL002/pJW007 pJW031; cm<sup>R</sup> kan<sup>R</sup> | This study |
| EcJW218 | EcDL002/pJW007 pJW032; cm<sup>R</sup> kan<sup>R</sup> | This study |
| EcJW219 | EcDL002/pJW007 pJW033; cm<sup>R</sup> kan<sup>R</sup> | This study |
| EcJW220 | EcDL002/pJW007 pJW034; cm<sup>R</sup> kan<sup>R</sup> | This study |
| EcJW221 | EcDL002/pJW007 pJW035; cm<sup>R</sup> kan<sup>R</sup> | This study |
| EcJW109 | EcDL002/pJW027; kan<sup>R</sup> | This study |
| EcJW110 | EcDL002/pJW028; kan<sup>R</sup> | This study |
| EcJW111 | EcDL002/pJW029; kan<sup>R</sup> | This study |
| EcJW112 | EcDL002/pJW030; kan<sup>R</sup> | This study |
| EcJW113 | EcDL002/pJW031; kan<sup>R</sup> | This study |
| EcJW114 | EcDL002/pJW032; kan<sup>R</sup> | This study |
| EcJW115 | EcDL002/pJW033; kan<sup>R</sup> | This study |
| EcJW116 | EcDL002/pJW034; kan<sup>R</sup> | This study |
| EcJW117 | EcDL002/pJW035; kan<sup>R</sup> | This study |

### Plasmid construction

The list of plasmids and primers used in this study are presented in Table 2 and Table 3, respectively. Pathway construction includes pyruvate-to-lactate ester modules and a library of upstream and downstream modules with various plasmid copy numbers, promoters, and ribosome binding sites (RBSs).

**Table 2.** A list of plasmids used in this study

| Plasmids | Genotypes | Sources |
| --- | --- | --- |
| pACYCDuet-1 | Two sets of MCS, T<sub>7</sub> promoter, P15A ori; cm<sup>R</sup> | Novagen |
| pETDuet-1 | Two sets of MCS, T<sub>7</sub> promoter, ColE1 ori; amp<sup>R</sup> | Novagen |
| pRSFDuet-1 | Two sets of MCS, T<sub>7</sub> promoter, RSF1030 ori; kan<sup>R</sup> | Novagen |
| pETite* | T<sub>7</sub> promoter, pBR322 ori; kan<sup>R</sup> | [10] |
| pCT24 | pETite* P<sub>T7</sub>::*pdC::adhB*::T<sub>T7</sub>; kan<sup>R</sup> | [10] |
| pCT13 | pCOLA P<sub>T7</sub>::*alsS::ilvC::ilvD*-P<sub>T7</sub>::*kivD::adhE*::T<sub>T7</sub>; kan<sup>R</sup> | [58] |
| pDL004 | pETite* *ATF1*; kan<sup>R</sup> | [13] |
| pDL005 | pETite* *ATF2*; kan<sup>R</sup> | [13] |
| pDL001 | pETite* *SAAT*; kan<sup>R</sup> | [13] |
| pDL006 | pETite* *VAAT*; kan<sup>R</sup> | [13] |
| pCT16 | pETite* *atfA*; kan<sup>R</sup> | [59] |
| pJW001 | pETite* P<sub>T7</sub>::*ldhA::pct*::T<sub>T7</sub>; amp<sup>R</sup> | This study |
| pJW002 | pJW001 P<sub>T7</sub>::*ldhA::pct*-P<sub>T7</sub>::*ATF1*::T<sub>T7</sub>; amp<sup>R</sup> | This study |
| pJW003 | pJW001 P<sub>T7</sub>::*ldhA::pct*-P<sub>T7</sub>::*ATF2*::T<sub>T7</sub>; amp<sup>R</sup> | This study |
| pJW004 | pJW001 P<sub>T7</sub>::*ldhA::pct*-P<sub>T7</sub>::*SAAT*::T<sub>T7</sub>; amp<sup>R</sup> | This study |
| pJW005 | pJW001 P<sub>T7</sub>::*ldhA::pct*-P<sub>T7</sub>::*VAAT*::T<sub>T7</sub>; amp<sup>R</sup> | This study |
| pJW006 | pJW001 P<sub>T7</sub>::*ldhA::pct*-P<sub>T7</sub>::*atfA*::T<sub>T7</sub>; amp<sup>R</sup> | This study |
| pJW007 | pACYCDuet-1 P<sub>T7</sub>::*ldhA::pdc::adhB*::T<sub>T7</sub>; cm<sup>R</sup> | This study |
| pJW008 | pETDuet-1 P<sub>T7</sub>::*ldhA::pdc::adhB*::T<sub>T7</sub>; amp<sup>R</sup> | This study |
| pJW009 | pRSFDuet-1 P<sub>T7</sub>::*ldhA::pdc::adhB*::T<sub>T7</sub>; kan<sup>R</sup> | This study |
| pJW010 | pACYCDuet-1 P<sub>T7</sub>::*pct::VAAT*::T<sub>T7</sub>; cm<sup>R</sup> | This study |
| pJW011 | pETDuet-1 P<sub>T7</sub>::*pct::VAAT*::T<sub>T7</sub>; amp<sup>R</sup> | This study |
| pJW012 | pRSFDuet-1 P<sub>T7</sub>::*pct::VAAT*::T<sub>T7</sub>; kan<sup>R</sup> | This study |
| pJW013 | pACYCDuet-1 P<sub>T7</sub>::*ldhA::pdc::adhB*-P<sub>T7</sub>::*pct::VAAT*::T<sub>T7</sub>; cm<sup>R</sup> | This study |
| pJW014 | pETDuet-1 P<sub>T7</sub>::*ldhA::pdc::adhB*-P<sub>T7</sub>::*pct::VAAT*::T<sub>T7</sub>; amp<sup>R</sup> | This study |
| pJW015 | pRSFDuet-1 P<sub>T7</sub>::*ldhA::pdc::adhB*-P<sub>T7</sub>::*pct::VAAT*::T<sub>T7</sub>; kan<sup>R</sup> | This study |
| pJW016 | pACYCDuet-1 P<sub>T7</sub>::*ldhA*::T<sub>T7</sub>-P<sub>T7</sub>::T<sub>T7</sub>; cm<sup>R</sup> | This study |
| pJW017 | pACYCDuet-1 P<sub>T7</sub>::*ldhA*::T<sub>T7</sub>-P<sub>AY1</sub>::T<sub>T7</sub>; cm<sup>R</sup> | This study |
| pJW018 | pACYCDuet-1 P<sub>T7</sub>::*ldhA*::T<sub>T7</sub>-P<sub>AY3</sub>::T<sub>T7</sub>; cm<sup>R</sup> | This study |
| pJW019 | pACYCDuet-1 P<sub>T7</sub>::*ldhA*::T<sub>T7</sub>-P<sub>AY1</sub>::synRBS<sub>pdC#1</sub>::*pdC::adhB*::T<sub>T7</sub>; cm<sup>R</sup> | This study |
| pJW020 | pACYCDuet-1 P<sub>T7</sub>::*ldhA*::T<sub>T7</sub>-P<sub>AY1</sub>::synRBS<sub>pdC#2</sub>::*pdC::adhB*::T<sub>T7</sub>; cm<sup>R</sup> | This study |
| pJW021 | pACYCDuet-1 P<sub>T7</sub>::*ldhA*::T<sub>T7</sub>-P<sub>AY3</sub>::synRBS<sub>pdC#3</sub>::*pdC::adhB*::T<sub>T7</sub>; cm<sup>R</sup> | This study |
| pJW022 | pACYCDuet-1 P<sub>T7</sub>::*ldhA*::T<sub>T7</sub>-P<sub>AY3</sub>::synRBS<sub>pdC#4</sub>::*pdC::adhB*::T<sub>T7</sub>; cm<sup>R</sup> | This study |
| pJW023 | pRSFDuet-1 P<sub>T7</sub>::*pct*::T<sub>T7</sub>-P<sub>T7</sub>::T<sub>T7</sub>; kan<sup>R</sup> | This study |
| pJW024 | pRSFDuet-1 P<sub>T7</sub>::synRBS<sub>pct#1</sub>::*pct*::T<sub>T7</sub>-P<sub>T7</sub>::T<sub>T7</sub>; kan<sup>R</sup> | This study |
| pJW025 | pRSFDuet-1 P<sub>T7</sub>::synRBS<sub>pct#2</sub>::*pct*::T<sub>T7</sub>-P<sub>T7</sub>::T<sub>T7</sub>; kan<sup>R</sup> | This study |
| pJW026 | pRSFDuet-1 P<sub>T7</sub>::synRBS<sub>pct#3</sub>::*pct*::T<sub>T7</sub>-P<sub>T7</sub>::T<sub>T7</sub>; kan<sup>R</sup> | This study |
| pJW027 | pRSFDuet-1 P<sub>T7</sub>::synRBS<sub>pct#1</sub>::*pct*::T<sub>T7</sub>-P<sub>T7</sub>::synRBS<sub>VAAT#1</sub>::*VAAT*::T<sub>T7</sub>; kan<sup>R</sup> | This study |
| pJW028 | pRSFDuet-1 P<sub>T7</sub>::synRBS<sub>pct#1</sub>::*pct*::T<sub>T7</sub>-P<sub>T7</sub>::synRBS<sub>VAAT#2</sub>::*VAAT*::T<sub>T7</sub>; kan<sup>R</sup> | This study |
| pJW029 | pRSFDuet-1 P<sub>T7</sub>::synRBS<sub>pct#1</sub>::*pct*::T<sub>T7</sub>-P<sub>T7</sub>::synRBS<sub>VAAT#3</sub>::*VAAT*::T<sub>T7</sub>; kan<sup>R</sup> | This study |
| pJW030 | pRSFDuet-1 P<sub>T7</sub>::synRBS<sub>pct#2</sub>::*pct*::T<sub>T7</sub>-P<sub>T7</sub>::synRBS<sub>VAAT#1</sub>::*VAAT*::T<sub>T7</sub>; kan<sup>R</sup> | This study |
| pJW031 | pRSFDuet-1 P<sub>T7</sub>::synRBS<sub>pct#2</sub>::*pct*::T<sub>T7</sub>-P<sub>T7</sub>::synRBS<sub>VAAT#2</sub>::*VAAT*::T<sub>T7</sub>; kan<sup>R</sup> | This study |
| pJW032 | pRSFDuet-1 P<sub>T7</sub>::synRBS<sub>pct#2</sub>::*pct*::T<sub>T7</sub>-P<sub>T7</sub>::synRBS<sub>VAAT#3</sub>::*VAAT*::T<sub>T7</sub>; kan<sup>R</sup> | This study |
| pJW033 | pRSFDuet-1 P<sub>T7</sub>::synRBS<sub>pct#3</sub>::*pct*::T<sub>T7</sub>-P<sub>T7</sub>::synRBS<sub>VAAT#1</sub>::*VAAT*::T<sub>T7</sub>; kan<sup>R</sup> | This study |
| pJW034 | pRSFDuet-1 P<sub>T7</sub>::synRBS<sub>pct#3</sub>::*pct*::T<sub>T7</sub>-P<sub>T7</sub>::synRBS<sub>VAAT#2</sub>::*VAAT*::T<sub>T7</sub>; kan<sup>R</sup> | This study |
| pJW035 | pRSFDuet-1 P<sub>T7</sub>::synRBS<sub>pct#3</sub>::*pct*::T<sub>T7</sub>-P<sub>T7</sub>::synRBS<sub>VAAT#3</sub>::*VAAT*::T<sub>T7</sub>; kan<sup>R</sup> | This study |

**Table 3.**
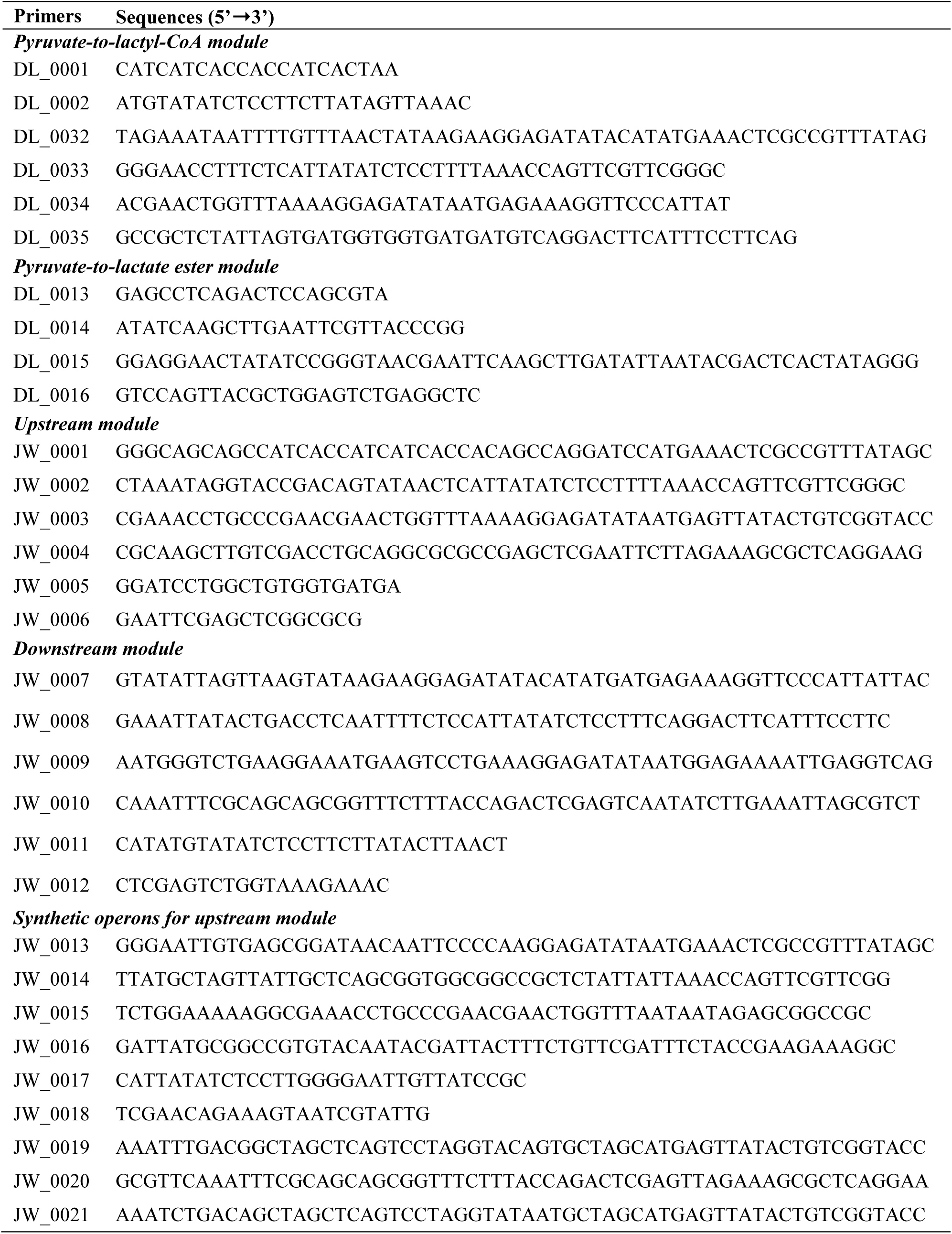
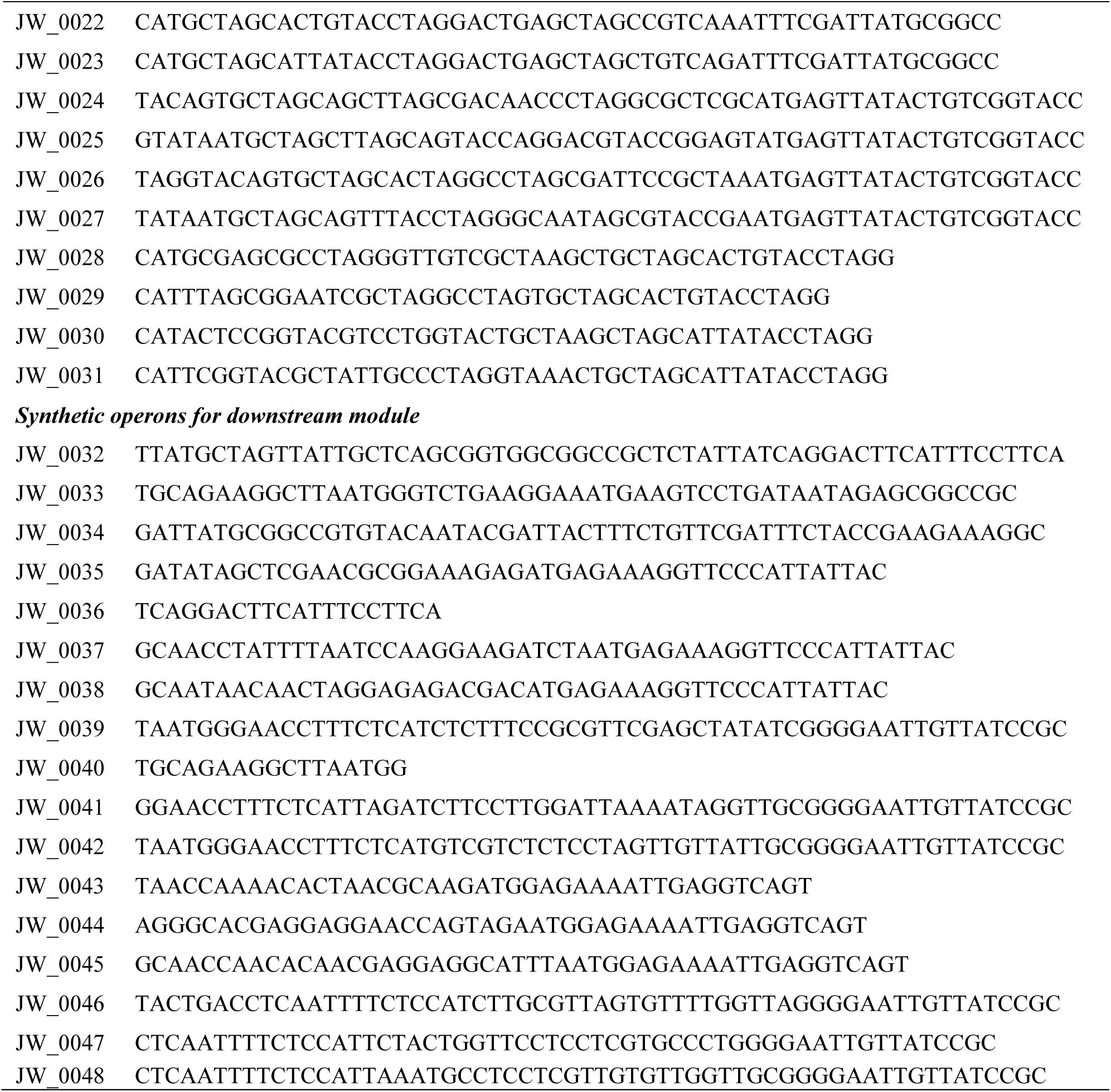
A list of primers used in this study

#### Construction of pyruvate-to-lactate ester modules

A library of pyruvate-to-lactate ester modules with five divergent AATs were constructed to screen for an efficient AAT for production of lactate esters via two rounds of cloning. First, the pyruvate-to-lactyl-CoA module (pJW001) was constructed by assembling three DNA fragments: i) the *ldhA* gene, encoding D-lactate dehydrogenase, amplified from *E. coli* MG1655 genomic DNA using the primer pair DL_0032/DL_0033, ii) the *pct* gene, encoding propionate CoA-transferase, amplified from *Clostridium propionicum* genomic DNA using the primer pair DL_0034/DL_0035, and iii) the backbone amplified from pETite* using the primer pair DL_0001/DL_0002 [49]. Then, the pyruvate-to-lactate ester modules (pJW002-006) were constructed by assembling three DNA fragments: i) the pyruvate-to-lactyl-CoA module amplified from pJW001 using the primer pair DL_0032/DL_0014, ii) the *ATF1* gene amplified from pDL004 for pJW002, the *ATF2* gene amplified from pDL005 for pJW003, the *SAAT* gene amplified from pDL001 for pJW004, the *VAAT* gene amplified from pDL006 for pJW005, or the *atfA* gene amplified from pCT16 for pJW006, using the primer pair DL_0015/DL_0016, and iii) the backbone amplified from pETite* using the primer pair DL_0013/ DL_0002. The genes *ATF1* and *ATF2* are originated from *Saccharomyces cerevisiae* [50], whereas the genes *SAAT*, *VAAT* and *atfA* are derived from *Fragaria ananassa* [51], *F. vesca* [52], and *Acinetobacter* sp. ADP1 [53], respectively.

#### Construction of a library of upstream and downstream modules with various plasmid copy numbers

A library of upstream and downstream modules were constructed to improve ethyl lactate biosynthesis through a combinatorial pathway optimization strategy using three different plasmids: i) pACYCDuet-1 (P15A origin of replication), ii) pETDuet-1 (ColE1 origin), and iii) pRSFDuet-1 (RSF1030 origin), having the plasmid copy numbers of 10, 40, and 100, respectively [54].

The upstream modules (pJW007-009) were constructed by assembling three DNA fragments: i) the *ldhA* gene amplified from pJW001 using the primer pair JW_0001/JW_0002, ii) the ethanol module containing *pdc* and *adhB* genes amplified from pCT24 using the primer pair JW_0003/JW_0004, and iii) the backbone amplified from pACYCDuet-1 for pJW007, from pETDuet-1 for pJW008, or from pRSFDuet-1 for pJW009 using the primer pair JW_0005/JW_0006.

The downstream modules (pJW010-012) were constructed by assembling three DNA fragments: i) the *pct* gene amplified from pJW001 using the primer pair JW_0007/JW_0008, ii) the *VAAT* gene amplified from pJW005 using the primer pair JW_0009/JW_0010, and iii) the backbone amplified from pACYCDuet-1 for pJW010, pETDuet-1 for pJW011, or pRSFDuet-1 for pJW012 using the primer pair JW_0011/JW_0012.

The combined upstream and downstream modules (pJW013-015) were constructed by assembling two DNA fragments: i) the upstream module amplified from pJW007 using the primer pair JW_0001/JW_0004 and ii) the backbone containing the downstream module amplified from pJW010 for pJW013, pJW011 for pJW014, or pJW012 for pJW015 using the primer pair JW_0005/JW_0006.

#### Construction of a library of upstream and downstream modules with various promoters and RBSs

For tighter regulation of biosynthetic pathway of ethyl lactate, we constructed the upstream and downstream modules with tunable promoters and RBSs.

The upstream modules (pJW019-022) were constructed via three rounds of cloning. First, the T7 terminator (T*_T7_*) was added between the multiple cloning site 1 (MCS1) and MCS2 of the pACYCDuet-1 backbone to create the first intermediate plasmid, pJW016, by assembling three DNA fragments: i) the *ldhA* gene amplified from pJW001 using the primer pair JW_0013/JW_0014, ii) the linker containing T*_T7_* terminator amplified from pETite* using the primer pair JW_0015/JW_0016, and iii) the backbone amplified from pACYCDuet-1 using the primer pair JW_0017/JW_0018. Next, the original T7 promoter (P*_T7_*) in MCS2 of pJW016 was replaced with the P*_AY1_* (BBa_J23100) promoter and P*_AY3_* (BBaJ23108) promoter to generate two second intermediate plasmids, pJW017 and pJW018, respectively, by assembling two DNA fragments: i) the ethanol module amplified from pCT24 under the P*_AY1_* promoter for pJW017 or P*_AY3_* promoter for pJW018 using the primer pair JW_0019/JW_0020 or JW_0021/JW_0020, respectively, and ii) the backbone amplified from pJW016 using the primer pair JW_0022/JW_0012 or JW_0023/JW_0012, respectively. Lastly, the final four synthetic operons (pJW019-022) were constructed by assembling two DNA fragments: i) the ethanol module amplified from pCT24 with the synthetic RBS sequences with predicted translation initiation rates of 0.33au for pJW019 and pJW021 and 0.03au for pJW020 and pJW022 using the primer pairs JW_0024/JW_0020, JW_0025/JW_0020, JW_0026/JW_0020, and JW_0027/JW_0020, respectively, and ii) the backbone amplified from pJW017 for pJW019, pJW017 for pJW020, pJW018 for pJW021, and pJW018 for pJW022 using the primer pairs JW_0028/JW_0012, JW_0029/JW_0012, JW_0030/JW_0012, and JW_0031/JW_0012, respectively. The P*_AY1_* and P*_AY3_* promoter sequences were obtained from the iGEM Anderson promoter library (http://parts.igem.org/Promoters/Catalog/Anderson) and the strength of promoters were assigned as P*_AY3_* = 0.5 x P*_AY1_*. The RBS Calculator v2.0 [55, 56] was used to generate four synthetic RBS sequences with predicted translation initiation rates of 0.33 and 0.03 between the P*_AY1_* (or P*_AY3_*) promoter and *pdc* start codon (Fig. S3).

The downstream modules (pJW027-035) were constructed via three rounds of cloning. First, the T*_T7_* terminator was added between MCS1 and MCS2 of the pRSFDuet-1 backbone to generate the first intermediate plasmid, pJW023, by assembling three DNA fragments: i) the *pct* gene amplified from pJW001 using the primer pair JW_0013/JW_0032, ii) the linker containing T*_T7_* terminator from pETite* using the primer pair JW_0033/JW_0034, and iii) the backbone from pRSFDuet-1 using the primer pair JW_0017/JW_0018. Then, the original RBS in MCS1 of pJW023 was replaced with synthetic RBSs of various strengths to generate the second intermediate plasmids, pJW024-026, by assembling two DNA fragments: i) the *pct* gene amplified from pJW001 with the synthetic RBS sequences with predicted translation initiation rates at 90, 9000, or 90000au for pJW024, pJW025 or pJW026 using the primer pair JW_0035/JW_0036, JW_0037/JW_0036, or JW_0038/JW_0036, respectively, and ii) the backbone amplified from pJW023 using the primer pair JW_0039/JW_0040 for pJW024, JW_0041/JW_0040 for pJW025, or JW_0042/JW_0040 for pJW026, respectively. Lastly, the final nine downstream modules (pJW027-035) were constructed by assembling a combination of two DNA fragments: i) the *VAAT* gene amplified from pDL006 with the synthetic RBS sequences predicted with translation initiation rates of 90, 9000, or 90000au for pJW027/pJW030/pJW033, pJW028/pJW031/pJW034, or pJW029/pJW032/pJW035 using the primer pair JW_0043/JW_0010, JW_0044/JW_0010, or JW_0045/JW_0010, respectively, and ii) the backbone amplified from pJW024, pJW025, or pJW026 for pJW027-029, pJW030-032, or pJW033-035 using the primer pair JW_0046/JW_0012, JW_0047/JW_0012 or JW_0048/JW_0012, respectively. The RBS Calculator v2.0 [55, 56] was used to generate six synthetic RBS sequences with predicted translation initiation rates of 90, 9000, and 90000au between the P*_T7_* promoter and *pct* (or *VAAT*) start codon (Fig. S3).

### Culture media and conditions

#### Culture media

For molecular cloning, seed cultures, and protein expression analysis, the Luria-Bertani (LB) medium, comprising of 10 g/L peptone, 5 g/L yeast extract, and 5 g/L NaCl, was used. For high-cell density cultures, pre-cultures of bioreactor batch fermentations, and growth inhibition analysis of lactate esters, the M9 hybrid medium [10] with 20 g/L glucose was used. For bioreactor batch fermentations, the M9 hybrid medium with 50 g/L glucose and 100 µL of antifoam (Antifoam 204, Sigma-Aldrich, MO, USA) was used. 30 µg/mL chloramphenicol (cm), 50 µg/mL kanamycin (kan), and/or 50 µg/mL ampicillin (amp) was added to the media for selection where applicable.

#### High-cell density cultures

For seed cultures, 2% (v/v) of stock cells were grown overnight in 5 mL of LB with appropriate antibiotics. For pre-cultures, 1% (v/v) of seed cultures were transferred into 100 mL of LB medium in 500 mL baffled flasks. For main cultures, pre-cultures were aerobically grown overnight (at 37°C, 200 rpm), centrifuged (4700 rpm, 10 min), and resuspended to yield an optical density measured at 600nm (OD_600nm_) of 3 in M9 hybrid medium containing appropriate concentration of isopropyl-beta-D-thiogalatopyranoside (IPTG) and antibiotics. The resuspended cultures were distributed into 15 mL polypropylene centrifuge tubes (Thermo Scientific, IL, USA) with a working volume of 5 mL and grown for 24 hour (h) on a 75° angled platform in a New Brunswick Excella E25 at 37°C, 200 rpm. The tubes were capped to generate anaerobic condition. All high-cell density culture studies were performed in biological triplicates.

#### pH-Controlled bioreactor batch fermentations

pH-Controlled bioreactor batch fermentations were conducted with a Biostat B+ (Sartorius Stedim, NY, USA) dual 1.5 L fermentation system at a working volume of 1 L M9 hybrid medium. The seed and pre-cultures were prepared as described in high-cell density cultures in LB and M9 hybrid media, respectively. For main cultures, 10% (v/v) of pre-cultures were inoculated into fermentation cultures. During the fermentation, to achieve high cell density, dual-phase fermentation approach [25, 57], aerobic cell growth phase followed by anaerobic production phase, was applied. For the first aerobic phase, the temperature, agitation, and air flow rate were maintained at 37°C, 1000 rpm, and 1 volume/volume/min (vvm) for 4 h, respectively. Then, the oxygen in the medium was purged by sparing nitrogen gas at 2 vvm for 2 h to generate anaerobic condition. For the subsequent anaerobic phase, 0.5 mM of IPTG was added to induce the protein expression, and the culture temperature and nitrogen flow rate were maintained at 30°C and 0.2 vvm, respectively. During the fermentation, the pH was maintained at 7.0 with 5 M KOH and 40% H_3_PO_4_. Bioreactor batch fermentation studies were performed in biological duplicates.

#### Growth inhibition analysis of lactate esters

Seed cultures of EcDL002 were prepared as described in high-cell density cultures. 4 % (v/v) of seed cultures were inoculated into 100 µL of the M9 hybrid media, containing various concentrations (0.5∼40 g/L) of lactate esters including ethyl-, propyl-, butyl-, isobutyl-, isoamyl-, or benzyl lactate, in a 96-well microplate. Then, the microplate was sealed with a plastic adhesive sealing film, SealPlate® (EXCEL Scientific, Inc., CA, USA) to prevent evaporation of lactate esters and incubated at 37°C with continuous shaking using a BioTek Synergy HT microplate reader (BioTek Instruments, Inc., VT, USA). OD_600nm_ was measured at 20 min intervals. Growth inhibition studies of lactate esters were performed twice in biological triplicates (*n* = 6).

### Protein expression and SDS-PAGE analysis

Seed cultures were prepared as described in high-cell density cultures. 1% (v/v) of seed cultures subsequently inoculated in 500 mL baffled flasks containing 100 ml of LB medium. Cells were aerobically grown at 37°C and 200 rpm and induced at an OD_600nm_ of 0.6∼0.8 with 0.5 mM of IPTG. After 4 h of induction, cells were collected by centrifugation and resuspended in 100 mM of sodium phosphate buffer (pH7.0) at the final OD_600nm_ of 10. Cell pellets were disrupted using a probe-type sonicator (Model 120, Fisher Scientific, NH, USA) on ice-water mixture. The resulting crude extracts were mixed with 6x sodium dodecyl sulfate (SDS) sample buffer, heated at 100°C for 5 min, and then analyzed by SDS-polyacrylamide gel electrophoresis (SDS-PAGE, 14% polyacrylamide gel). Protein bands were visualized with Coomassie Brilliant Blue staining.

### Analytical methods

#### Determination of cell concentrations

The optical density was measured at 600 nm using a spectrophotometer (GENESYS 30, Thermo Scientific, IL, USA). The dry cell mass was obtained by multiplication of the optical density of culture broth with a pre-determined conversion factor, 0.48 g/L/OD.

#### High performance liquid chromatography (HPLC)

Glucose, lactate, acetate, ethanol, isobutanol, isoamyl alcohol, and benzyl alcohol were quantified by using the Shimadzu HPLC system (Shimadzu Inc., MD, USA) equipped with the Aminex HPX-87H cation exchange column (BioRad Inc., CA, USA) heated at 50°C. A mobile phase of 10 mN H_2_SO_4_ was used at a flow rate of 0.6 mL/min. Detection was made with the reflective index detector (RID) and UV detector (UVD) at 220 nm.

#### Gas chromatography coupled with mass spectroscopy (GC/MS)

All esters were quantified by GC/MS. For GC/MS analysis, analytes in the supernatants were extracted with dichloromethane (DCM), containing pentanol as an internal standard, in a 1:1 (v/v) ratio for 1 h at 37°C, 200 rpm in 15 mL polypropylene centrifuge tubes. After extraction, supernatant-DCM mixtures were centrifuged and 5 μL of DCM extracts were injected into a gas chromatograph (GC) HP 6890 equipped with the mass selective detector (MS) HP 5973. For the GC system, helium was used as the carrier gas at a flow rate of 0.5 mL/min and the analytes were separated on a Phenomenex ZB-5 capillary column (30 m x 0.25 mm x 0.25 μm). The oven temperature was programmed with an initial temperature of 50°C with a 1°C/min ramp up to 58°C. Next a 25°C/min ramp was deployed to 235°C and then finally held a temperature of 300°C for 2 minutes to elute any residual non-desired analytes. The injection was performed using a splitless mode with an initial injector temperature of 280°C. For the MS system, a selected ion monitoring (SIM) mode was deployed to detect analytes.

The SIM parameters for detecting lactate esters were as follows: i) for pentanol, ions 53.00, 60.00, and 69.00 detected from 5.00 to 7.70 min, ii) for ethyl lactate, ions 46.00, 47.00, and 75.00 detected from 7.70 to 10.10 min, iii) for propyl lactate, ions 59.00, 88.00, and 89.00 detected from 10.10 to 11.00 min, iv) for isobutyl lactate, ions 56.00, 57.00, and 59.00 detected from 11.00 to 11.60 min, v) for butyl lactate, ions 75.00, 91.00, and 101.00 detected from 11.60 to 12.30 min, vi) for isoamyl lactate, ions 46.00, 73.00, 75.00 from 12.30 to 14.50 min, and vii) for benzyl lactate, ions 45.00, 91.00, and 180.00 from 14.50 to 15.08 min. The SIM parameters for detecting acetate esters were as follows: i) for ethyl acetate, ions 45.00, 61.00, and 70.00 detected from 4.22 to 5.35 min, ii) for propyl acetate, ions 57.00, 59.00, and 73.00 detected from 5.35 to 6.40 min, iii) for pentanol, ions 53.00, 60.00, and 69.00 detected from 6.40 to 6.60 min, iv) for isobutyl acetate, ions 56.00, 61.00, and 73.00 detected from 6.60 to 7.70 min, v) for butyl acetate, ions 57.00, 71.00, and 87.00 detected from 7.70 to 9.45 min, vi) for isoamyl acetate, ions 58.00, 70.00, and 88.00 detected from 9.45 to 13.10 min, and vii) for benzyl acetate, ions 63.00, 107.00, and 150.00 from 13.10 to 15.82 min.

#### Statistics

Statistical analysis was performed with SigmaPlot v.14 using the two-tailed unpaired Student’s t-test.

## ABBREVIATIONS

LdhA: lactate dehydrogenase
Pct: propionate CoA-transferase
AAT: alcohol acyltransferase
ATF1: alcohol acyltransferase from *Saccharomyces cerevisiae*
ATF2: alcohol acyltransferase from *Saccharomyces cerevisiae*
SAAT: alcohol acyltransferase from *Fragaria ananassa*
VAAT: alcohol acyltransferase from *Fragaria vesca*
AtfA: alcohol acyltransferase from *Acinetobacter* sp. ADP1
OD: optical density
DCW: dry cell weight
SDS-PAGE: sodium dodecyl sulfate-polyacrylamide gel electrophoresis
IPTG: isopropyl β-D-thiogalactopyranoside
MCS: multi cloning site
RBS: ribosome binding site
au: arbitrary unit
HPLC: high-performance liquid chromatography
GC/MS: gas chromatography coupled with mass spectrometry
SIM: selected ion monitoring
DCM: dichloromethane
rpm: revolutions per minute
v/v: volume per volume
vvm: volume per volume per minute.

## AUTHOR’S CONTRIBUTIONS

CTT conceived and supervised this study. JWL and CTT designed the experiments, analyzed the data, and drafted the manuscript. JWL performed the experiments. Both authors read and approved the final manuscript.

## ACKNOWLEDGMENTS

The authors would like to thank the Center of Environmental Biotechnology at UTK for using the GC/MS instrument.

## COMPETING INTERESTS

The authors declare that they have no competing interests.

## AVAILABILITY OF SUPPORTING DATA

Additional files 1 and 2 contain supporting data

## CONSENT FOR PUBLICATION

All the authors consent for publication.

## ETHICAL APPROVAL AND CONSENT TO PARTICIPATE

Not applicable

## FUNDING

This research was financially supported in part by the NSF CAREER award (NSF#1553250) and both the BioEnergy Science Center (BESC) and Center for Bioenergy Innovation (CBI), the U.S. Department of Energy (DOE) Bioenergy Research Centers funded by the Office of Biological and Environmental Research in the DOE Office of Science.

## Notes

#### Summary of Updates

The title is updated.

