## Supplementary Figures for "Microbial Biosynthesis of Lactate Esters"

Cong T. Trinh

Dept of Chemical and Biomolecular Engineering

University of Tennessee, Knoxville

1512 Middle Dr., DO#432

Knoxville, TN 37996

Additional Contact Information: Jong-Won Lee:

**Supplementary Figure S1.** Expression of the recombinant enzymes in engineered *E. coli* strains. The positions corresponding to the overexpressed proteins are indicated by arrowheads. Lane M represents protein ladder while lanes T, S, and I are referred to total, soluble, and insoluble proteins, respectively. ①~③, Pyruvate-to-lactate ester module; ④~⑤, Ethanol module; ⑥~⑩, Isobutanol module. Protein sizes were predicted with their amino acids sequences.

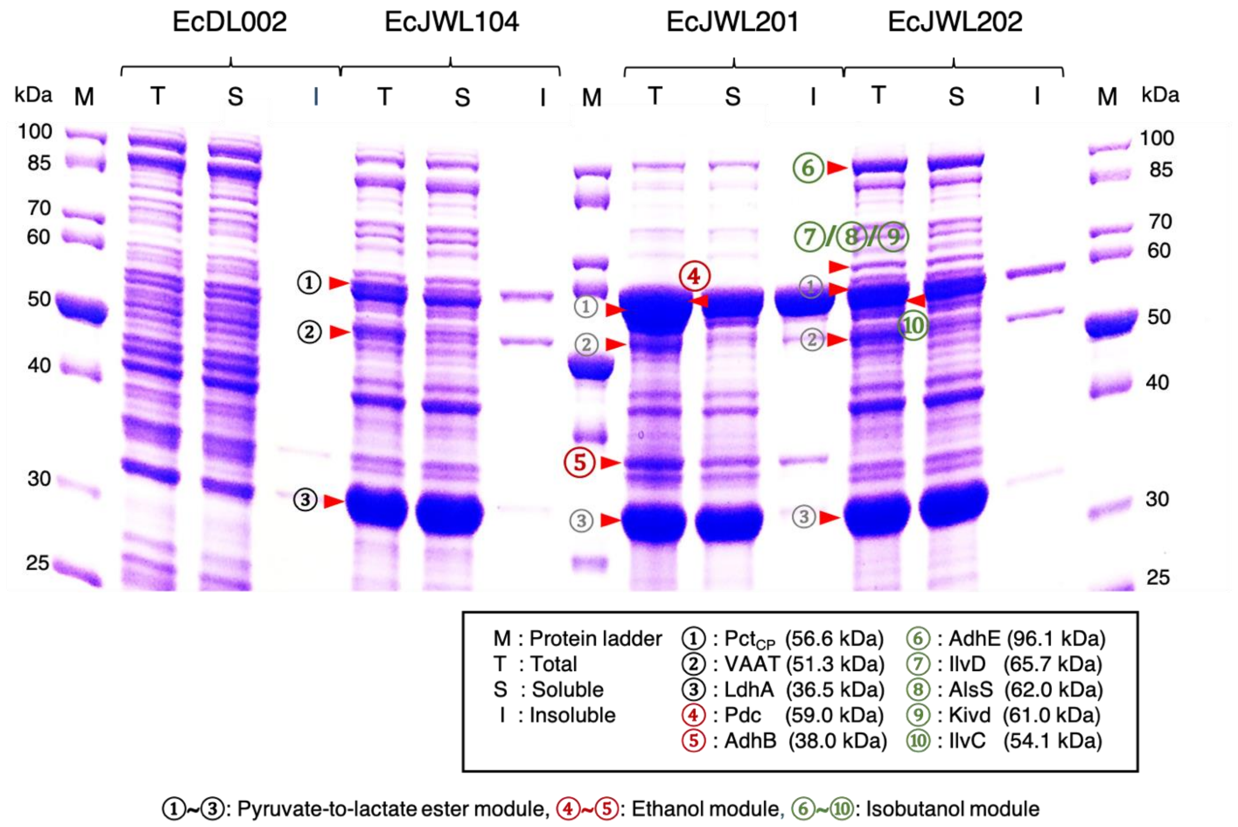

**Supplementary Figure S2.** Effect of lactate esters on cell growth. **(A)** Specific growth rates of EcDL002 with or without addition of lactate esters. (B) logP values of characterized lactate esters. The values were obtained from http://www.thegoodscentscompany.com. **(C-H)** Growth curves of EcDL002 with or without addition of **(C)** *n*-ethyl lactate (NEL), **(D)** *n*-propyl lactate (NPL), **(E)** *n*-butyl lactate (NBL), **(F)** *i*-butyl lactate (IBL), **(G)** *i*-amyl lactate (IAL), and **(H)** benzyl lactate (BZL).

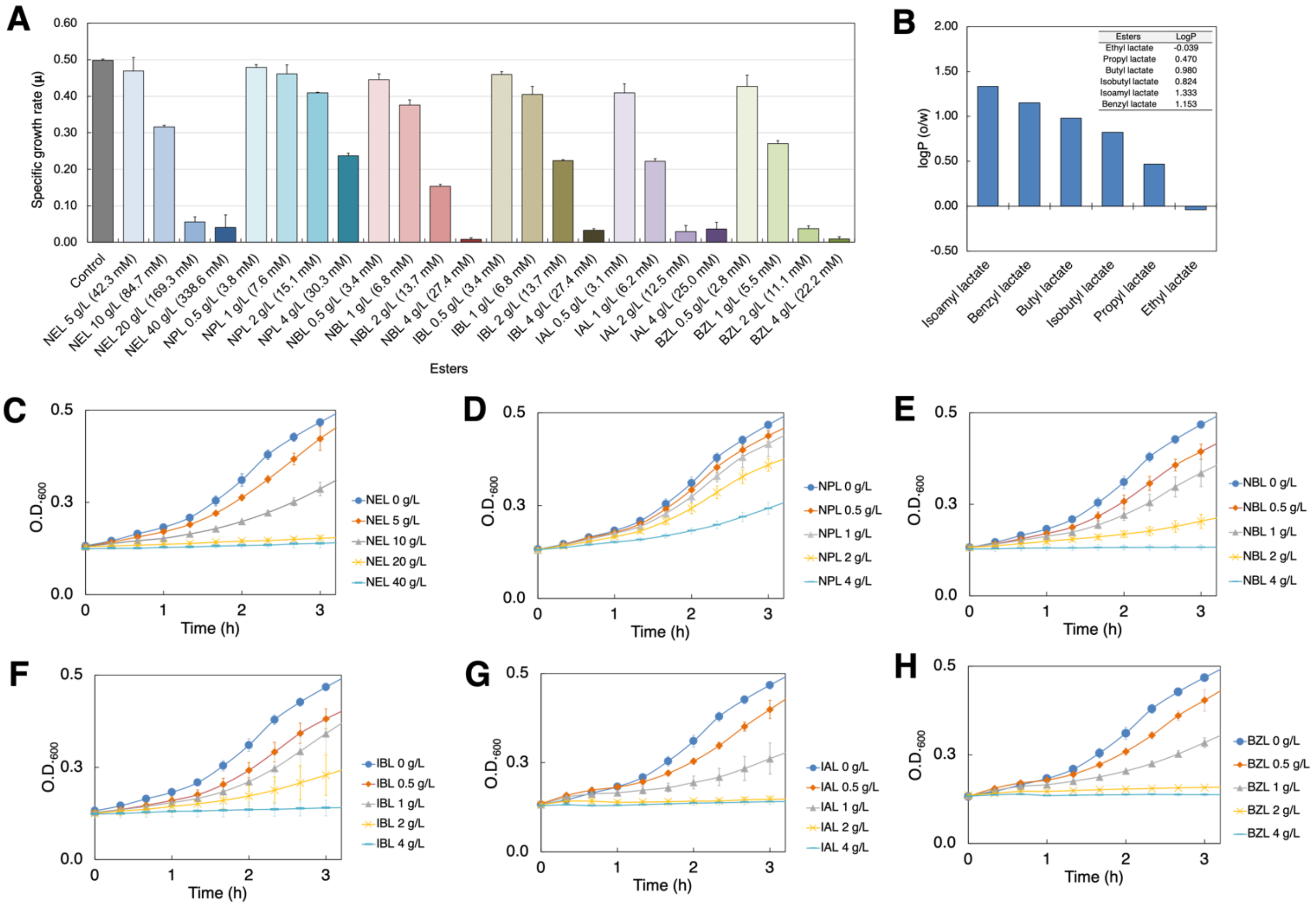

**Supplementary Figure S3.** Design of **(A)** upstream module and **(B)** downstream module of the ethyl lactate pathway. The RBS Calculator v2.0 software was used to generate synthetic RBS sequences. For the upstream, four synthetic RBS sequences were generated with predicted translation initiation rates at 0.33 and 0.03 between the $P_{AY1}$ or $P_{AY3}$ promoter and *pdc* start codon. For the downstream, six synthetic RBS sequences were generated with predicted translation initiation rates at 90, 9000, and 90000au between the $P_{T7}$ promoter and *pct* or *VAAT* start codon.

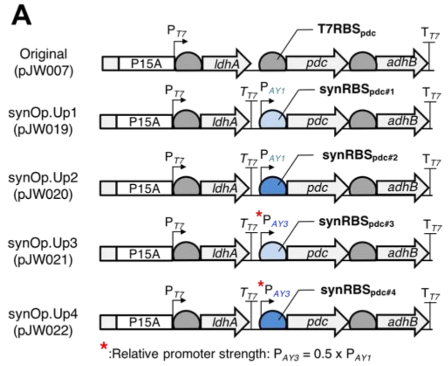


| RBS ID | Sequences | Predicted translation initiation rate (au) | Relative translation initiation rate (au) | Length (bp) |
|---|---|---|---|---|
| **$T7RBS_{pdc}$** | AAGGAGATATA | 33.11 | 1 | 11 |
| **$synRBS_{pdc\#1}$** | TTAGCAGTACCAGGACGTACCGGAGT | 0.33 | $10^{-2}$ | 27 |
| **$synRBS_{pdc\#2}$** | AGTTTACCTAGGGCAATAGCGTACCGA | 0.03 | $10^{-3}$ | 27 |
| **$synRBS_{pdc\#3}$** | AGCTTAGCGACAACCCTAGGCGCTCGC | 0.35 | $10^{-2}$ | 27 |
| **$synRBS_{pdc\#4}$** | ACTAGGCCTAGCGATTCCGCTAA | 0.03 | $10^{-3}$ | 23 |

$P_{AY1}$ (BBa_J23100) and $P_{AY3}$ (BBa_J23108) were obtained from iGEM Andersen promoter library

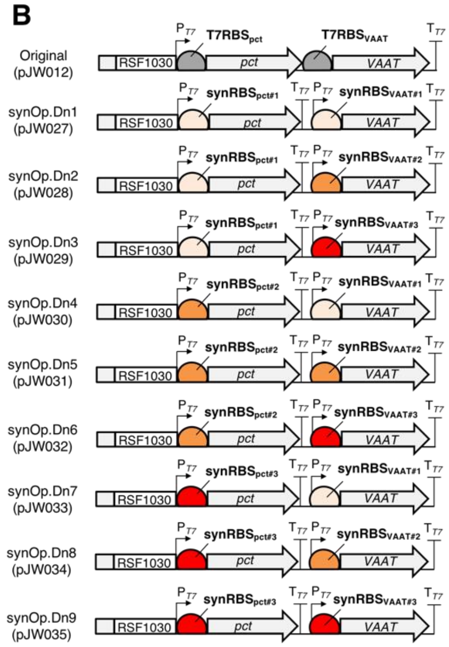


| RBS ID | Sequences | Predicted translation initiation rate (au) | Relative translation initiation rate (au) | Length (bp) |
|---|---|---|---|---|
| **$T7RBS_{pct}$** | AAGGAGATATA | 9417.53 | $10^{2}$ | 11 |
| **$synRBS_{pct\#1}$** | GATATAGCTCGAACGCGGAAAGAG | 84.56 | $10^{0}$ | 24 |
| **$synRBS_{pct\#2}$** | GCAACCTATTTTAATCCAAGGAAGATCT A | 8695.18 | $10^{2}$ | 29 |
| **$synRBS_{pct\#3}$** | GCAATAACAACTAGGAGAGACGAC | 95382.72 | $10^{3}$ | 24 |
| **$T7RBS_{VAAT}$** | AAGGAGATATA | 87.01 | 1 | 11 |
| **$synRBS_{VAAT\#1}$** | TAACCAAAACACTAACGCAAG | 93.02 | $10^{0}$ | 22 |
| **$synRBS_{VAAT\#2}$** | AGGGCACGAGGAGGAACCAGTAGA | 8228.14 | $10^{2}$ | 24 |
| **$synRBS_{VAAT\#3}$** | GCAACCAACACAACGAGGAGGCATTTA | 93714.75 | $10^{3}$ | 27 |

**Supplementary Figure S4. (A)** Correlation between ester production and the amount of added ethanol in high cell density cultures of EcJW209-212. **(B)** Correlation between ester production and the RBS strength for VAAT expression in high cell density culture of EcJW213-221.

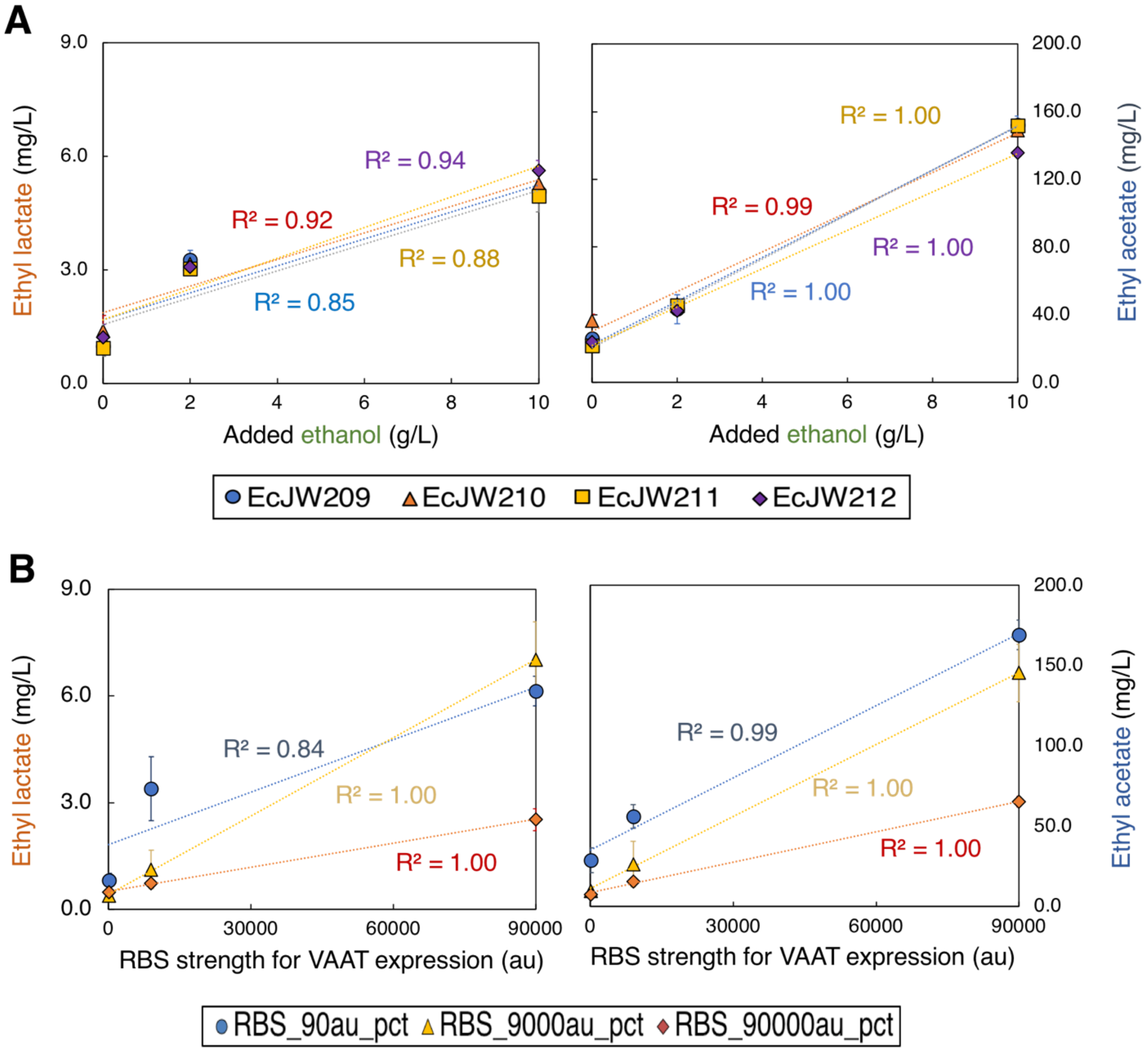
